# Cryo-EM structures of human RNA polymerase I

**DOI:** 10.1101/2021.05.31.446457

**Authors:** Agata D. Misiaszek, Mathias Girbig, Helga Grötsch, Florence Baudin, Aleix Lafita, Brice Murciano, Christoph W. Müller

## Abstract

RNA polymerase I (Pol I) specifically synthesizes ribosomal RNA. Pol I upregulation is linked to cancer, while mutations in the Pol I machinery lead to developmental disorders. Here, we report the cryo-EM structure of elongating human Pol I at 2.7 Å resolution. In the exit tunnel, we observe a double-stranded RNA helix that may support Pol I processivity. Our structure confirms that human Pol I consists of 13 subunits with only one subunit forming the Pol I stalk. Additionally, the structure of human Pol I in complex with the initiation factor RRN3 at 3.1 Å resolution reveals stalk flipping upon RRN3 binding. We also observe an inactivated state of human Pol I bound to an open DNA scaffold at 3.3 Å resolution. Lastly, the high-resolution structure of human Pol I allows mapping of disease-related mutations that can aid understanding of disease etiology.

## Introduction

RNA polymerase I (Pol I) is one of three eukaryotic RNA polymerases and is specialized in the transcription of ribosomal RNA (rRNA). In humans, Pol I exclusively transcribes the 43 kb rDNA gene. The produced 47S pre-rRNA transcript is later processed into 18S, 5.8S and 28S rRNAs which together with 5S rRNA produced by Pol III build the RNA core of the ribosome^1^. rRNA constitutes 80-90% of the total RNA mass in mammalian cells^2^. Owing to the high-energy expense imposed by rRNA transcription, Pol I activity needs to be tightly regulated.

Pol I transcription is the first step in ribosome biogenesis and thus plays a key role in cellular homeostasis^3^. Any misregulation of Pol I transcription can have deleterious effects including, for example, cancer. Upregulation of Pol I activity is required for cancer cells to proliferate. Therefore, Pol I transcription is a promising drug target for a large range of cancer types^4^. Several drugs acting on Pol I co-factors are already in clinical use, while other small molecules directly targeting Pol I are in clinical trials^3^. Transcriptional activity at the rDNA loci may compromise gene integrity, which promotes ageing^5^. Concomitantly, partial inhibition of Pol I has been associated with increasing longevity^6^. High rates of rRNA transcription are also observed in pluripotent stem cells^5^ and any impairment in the function of Pol I or its associated factors during development can lead to serious diseases resulting from impaired ribosome biogenesis and function, collectively named ribosomopathies^7^. Mutations causing Acrofacial Dysostosis (AD) have been mapped to the largest subunit of Pol I^8^, while some mutations causing Treacher Collins syndrome (TCS) have been found in the second largest Pol I subunit^9^. Mutations causing TCS are also found in the subunits shared between Pol I and Pol III^10,11^ making it difficult to distinguish whether functional impairment of Pol I or Pol III is causing the disease. The same difficulty applies to mutations associated with Hypomyelinating Leukodystrophy (HLD)^12^. A detailed understanding of the Pol I structure and function in humans is therefore required to better understand its role in development, ageing and in the etiology of complex diseases.

Pol I shares its general architecture with other eukaryotic DNA-dependent RNA polymerases which comprise of a homologous core bound by the stalk sub-complex^13^. In addition, Pol I and Pol III have stably integrated subunits homologous to general transcription factors of Pol II, namely TFIIE and TFIIF^14^. Pol I is the second largest among the eukaryotic RNA polymerases, both in terms of size and the number of subunits, with Pol II consisting of 12 subunits^15,16^ and Pol III comprising 17 subunits^17,18^. While yeast Pol I comprises 14 subunits^19,20^, in humans only homologs of 13 subunits have been identified^21^.

For transcription to take place, Pol I needs to bind to the promoter sequence in the context of accessory factors which form a pre-initiation complex (PIC). Pol I bound by RRN3 associates with the 5-subunit Selectivity Factor 1 (SL1) containing the TATA-box Binding Protein (TBP), which is activated by the Upstream Binding Factor (UBF) to allow specific rRNA transcription initiation^21^. All components of the human Pol I PIC are regulated by post-translational modifications to regulate the rate of rRNA production in response to cell cycle, growth factors, nutrient availability or stress^22–27^.

Past studies have provided a reasonable understanding of the structure and transcription cycle of yeast Pol I^19,20,28–35^. However, to understand the role of Pol I related pathologies, obtaining structural insights into human Pol I is necessary. Here, we report cryo-electron microscopy (cryo-EM) structures of human Pol I bound to different nucleic acid scaffolds as well as in complex with initiation factor RRN3 at 2.7 to 3.3 Å resolution. Insights drawn from these results expand our understanding of human Pol I function and lay the ground for further studies of its regulation and role in disease.

## Results

### Structure of elongating human Pol I

For cryo-EM structure determination, we purified human Pol I from a HEK293T suspension cell line where subunit RPAC1 was endogenously tagged by CRISPR-Cas9^17^. The introduced tag carries the mCherry fluorescent protein, which allowed for the confirmation of the correct cellular localization of Pol I in the nucleoli of the engineered cell line (Fig. 1a). The quality of the purified Pol I was assessed by SDS-PAGE, and the identity of all subunits was confirmed by mass spectrometry (Fig. 1b, Supplementary Table 1). Additionally, we confirmed the transcriptional activity of the obtained human Pol I through an *in vitro* RNA primer extension assay (Fig. 1c). Subsequently, we used the DNA-RNA scaffold, mimicking the elongation transcription bubble, for the cryo-EM structure determination of the Pol I elongating complex (Pol I EC). The human Pol I EC was resolved at 2.7 Å resolution with the local resolution in the center of the complex reaching up to 2.6 Å (Extended Data Fig. 1). To resolve the more flexible, peripheral parts of the complex, we used focused classification and multi-body refinement, which yielded three additional partial cryo-EM maps (Extended Data Fig. 1, Table. 1).

**Fig. 1:**
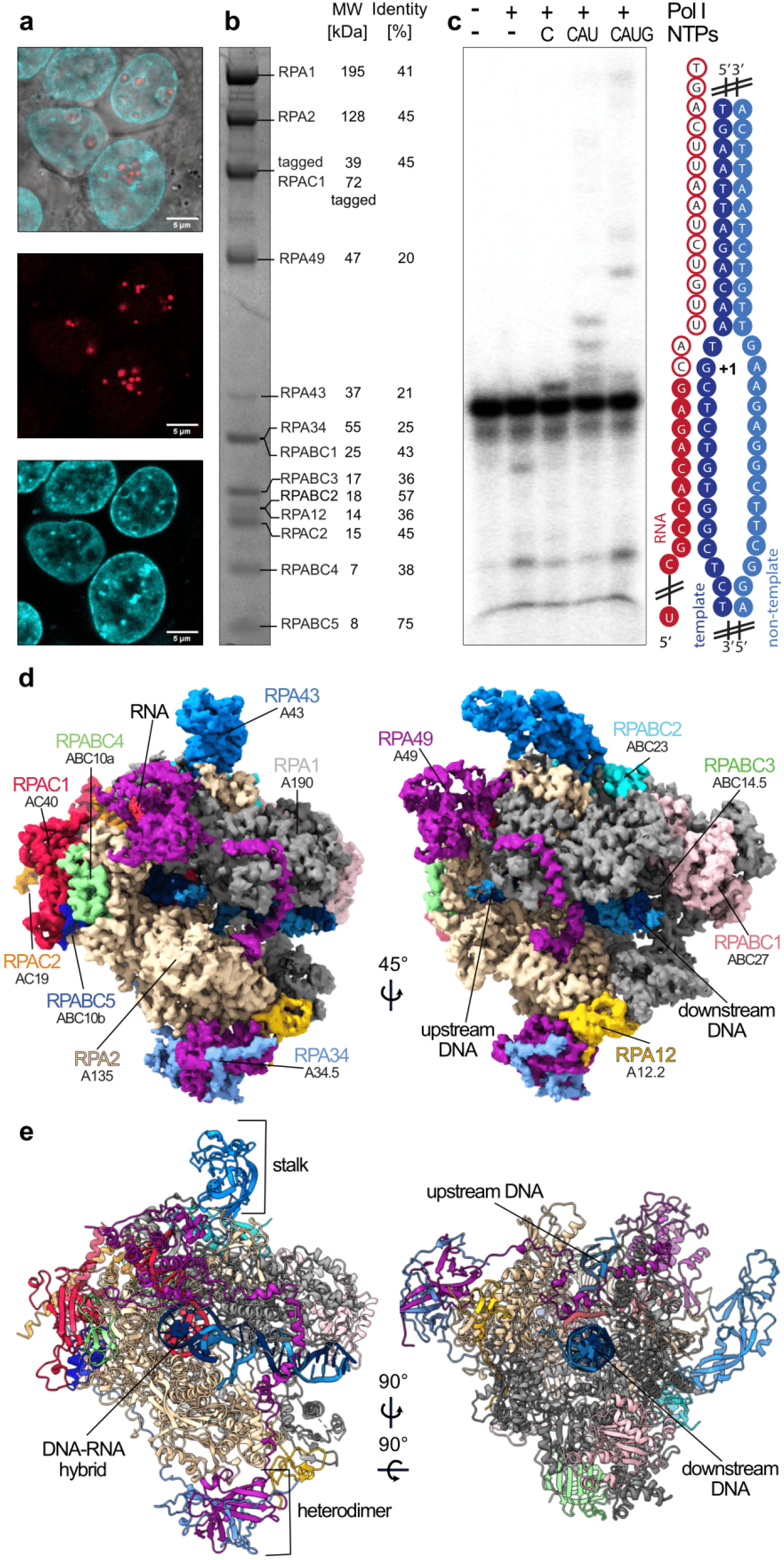
Structure of human Pol I. **a** Confocal images of the HEK293T cell line with endogenously tagged subunit RPAC1 showing localization to the nucleolus. Top panel: overlay of the transmission image with the mCherry signal (red, middle panel) and live Hoechst dye staining the DNA (cyan, bottom panel). Scale bar: 5 μm. **b** Coomassie-stained SDS-PAGE of the purified human Pol I. Identities of the bands labelled were confirmed by mass spectrometry (for details see Table S1). The percent sequence identity between yeast and human is shown on the right. **c** *In vitro* primer extension assay confirming the activity of the purified Pol I. 5’ radioactively tagged 19 base RNA primer (red) was assembled with the DNA scaffold (blue) with an artificial open bubble introduced by a mismatch (cartoon representation). Purified Pol I was incubated with the nucleic acid scaffold in presence of NTPs indicated at the top of the gel. For experimental details, see Methods. **d** Cryo-EM map of the Pol I EC (composite Map C in Table 1) colored according to its subunit composition. Subunits are labelled with the human nomenclature and (below) with the yeast counterpart. **e** Structural model of the Pol I EC with the functional subdomains (stalk and heterodimer) indicated.

**Table 1.** Cryo-EM data collection, refinement and validation statistics.

|  | Map A<br>(EMDB-12795)<br>EC Pol I<br>(PDB 7OB9) | Map B<br>(EMDB-12795)<br>Stalk tWH<br>Focused<br>Classification | Map B1<br>(EMDB-12795)<br>Multibody<br>Refinement<br>Upper Clamp | Map B2<br>(EMDB-12795)<br>Multibody<br>Refinement<br>Core | Map C<br>(EMDB-12795)<br>Composite |
| --- | --- | --- | --- | --- | --- |
| **Data collection and processing** |  |  |  |  |  |
| Magnification | 105,000 | 105,000 | 105,000 | 105,000 | 105,000 |
| Voltage (kV) | 300 | 300 | 300 | 300 | 300 |
| Electron exposure (e-/Å<sup>2</sup>) | 50.9 | 50.9 | 50.9 | 50.9 | 50.9 |
| Defocus range (µm) | 1.0-2.5 | 1.0-2.5 | 1.0-2.5 | 1.0-2.5 | 1.0-2.5 |
| Pixel size (Å) | 0.822 | 0.822 | 0.822 | 0.822 | 0.822 |
| Symmetry imposed | C1 | C1 | C1 | C1 | C1 |
| Initial particle images (no.) | 2,171,736 | 2,171,736 | 2,171,736 | 2,171,736 | 2,171,736 |
| Final particle images (no.) | 198,822 | 37,180 | 37,180 | 37,180 | - |
| Map resolution (Å) | 2.7 | 3.0 | 3.1 | 3.0 | - |
| FSC threshold | 0.143 | 0.143 | 0.143 | 0.143 | - |
| Map resolution range (Å) | 2.55-4.50 | 3.03-5.62 | 3.13-5.81 | 3.04-5.10 | - |
| **Refinement** |  |  |  |  |  |
| Initial model used<br>(PDB code) | 4C3I, 7AE1,<br>6LHR,<br>5M5X, 6RQT |  |  |  |  |
| Model resolution (Å) | 3.30 |  |  |  |  |
| Map sharpening *B* factor (Å<sup>2</sup>) | -67 |  |  |  |  |
| **Model composition** |  |  |  |  |  |
| Non-hydrogen atoms | 36471 |  |  |  |  |
| Protein residues | 4412 |  |  |  |  |
| Nucleotide residues | 69 |  |  |  |  |
| Ligands | 6x Zn, 1x Mg |  |  |  |  |
| ***B* factors (Å<sup>2</sup>)** |  |  |  |  |  |
| Protein | 109.26 |  |  |  |  |
| Ligand | 115.09 |  |  |  |  |
| **R.m.s. deviations** |  |  |  |  |  |
| Bond lengths (Å) | 0.008 |  |  |  |  |
| Bond angles (°) | 1.26 |  |  |  |  |
| **Validation** |  |  |  |  |  |
| MolProbity score | 1.60 |  |  |  |  |
| Clashscore | 4.64 |  |  |  |  |
| Poor rotamers (%) | 0.31 |  |  |  |  |
| **Ramachandran plot** |  |  |  |  |  |
| Favored (%) | 94.70 |  |  |  |  |
| Allowed (%) | 5.25 |  |  |  |  |
| Disallowed (%) | 0.05 |  |  |  |  |

**Table 1 (continued)**
|  | Map D<br>(EMDB-12796)<br>Pol I – RRN3 | Map E<br>(EMDB-12796)<br>Pol I – RRN3<br>Focused<br>Refinement<br>(PDB 7OBA) | Map F<br>(EMDB-12797)<br>Pol I OC<br>(PDB 7OBB) | Map G<br>(EMDB-12797)<br>Pol I OC<br>Nucleic acid<br>Focused<br>Classification |
| --- | --- | --- | --- | --- |
| **Data collection and processing** |  |  |  |  |
| Magnification | 130,000 | 130,000 | 130,000 | 130,000 |
| Voltage (kV) | 300 | 300 | 300 | 300 |
| Electron exposure (e-/Å<sup>2</sup>) | 41.26 | 41.26 | 41.26 | 41.26 |
| Defocus range (µm) | 0.5-2.25 | 0.5-2.25 | 0.5-2.25 | 0.5-2.25 |
| Pixel size (Å) | 0.822 | 0.822 | 0.822 | 0.822 |
| Symmetry imposed | C1 | C1 | C1 | C1 |
| Initial particle images (no.) | 2,628,144 | 2,628,144 | 2,628,144 | 2,628,144 |
| Final particle images (no.) | 169,513 | 260,363 | 175,912 | 164,436 |
| Map resolution (Å) | 3.2 | 3.1 | 3.3 | 3.3 |
| FSC threshold | 0.143 | 0.143 | 0.143 | 0.143 |
| Map resolution range (Å) | 3.0-7.0 | 2.9-7.3 | 3.0-5.9 | 3.0-6.7 |
| **Refinement** |  |  |  |  |
| Initial model used<br>(PDB code) |  | 4C3I, 7AE1 | 4C3I, 7AE1,<br>5M5W, 6RUO |  |
| Model resolution (Å) |  | 3.1 | 3.6 |  |
| Map sharpening *B* factor (Å<sup>2</sup>) |  | -103 | -91 |  |
| **Model composition** |  |  |  |  |
| Non-hydrogen atoms |  | 35684 | 33051 |  |
| Protein residues |  | 4399 | 4113 |  |
| Nucleotide residues |  | - | 17 |  |
| Ligands |  | 7x Zn | 7x Zn |  |
| ***B* factors (Å<sup>2</sup>)** |  |  |  |  |
| Protein |  | 65.99 | 75.19 |  |
| Ligand |  | 88.12 | 119.54 |  |
| **R.m.s. deviations** |  |  |  |  |
| Bond lengths (Å) |  | 0.0111 | 0.0097 |  |
| Bond angles (°) |  | 141 | 1.37 |  |
| **Validation** |  |  |  |  |
| MolProbity score |  | 1.77 | 1.76 |  |
| Clashscore |  | 5.33 | 5.21 |  |
| Poor rotamers (%) |  | 0.41 | 0.50 |  |
| **Ramachandran plot** |  |  |  |  |
| Favored (%) |  | 91.96 | 92.24 |  |
| Allowed (%) |  | 7.99 | 7.74 |  |
| Disallowed (%) |  | 0.05 | 0.02 |  |

We were able to assign cryo-EM densities to 13 Pol I subunits (Fig. 1d) and build the atomic model of the human Pol I EC (Fig. 1e). The general architecture consists of a horseshoe-shaped core bound by a stalk formed by the RPA43 subunit, resembling yeast Pol I. The core is further complemented by a TFIIE/F-like heterodimer, which consists of RPA34 and RPA49^36^. The C-terminus of RPA49 harbors a TFIIE-like tandem-winged-helix (tWH) domain^13,36,37^ that is observed in close proximity to the RNA exit tunnel (Fig. 1d, e). The well-resolved DNA-RNA scaffold enabled us to build a large portion of the downstream double-stranded DNA, the DNA-RNA hybrid in the active center, the RNA in the exit tunnel, as well as a portion of the upstream DNA (Fig. 1d, e). Overall, the general architecture of Pol I is conserved between yeast and humans despite the low sequence identity ranging from 20 to 45% for Pol I-specific subunits (Fig. 1b).

### Double-stranded RNA in the exit tunnel

For the assembly of the DNA-RNA scaffold mimicking the elongation bubble we used a 19-nuclotide RNA primer and a 43 bp DNA duplex containing a mismatch over 12 nucleotides. In the active site, we confidently assigned positions of all RNA bases in the DNA-RNA hybrid. Further, we could trace the RNA backbone extending into the RNA exit tunnel over a length of 16 nucleotides. This is remarkably different in comparison to similar structures of RNA polymerases in elongating states. In the structures of elongating human Pol III, only 5^38^ to 6^17^ nucleotides were visible, while in yeast Pol I between 7 and 13 nucleotides could be traced^28^. To our surprise, cryo-EM density bearing features of double-stranded RNA could be observed in the exit tunnel (Fig. 2a). The 5’ end of the RNA primer used is self-complementary to the exiting RNA strand (Fig. 2b), and thus we placed an ideal double-stranded RNA helix (A-form) into the exit tunnel that fit well into the cryo-EM density and remained intact after refinement (Fig. 2a). So far, RNA secondary structure elements in the exit tunnel of RNA polymerases have only been captured in bacterial RNA polymerase^39^. In human Pol I, the RNA exit tunnel is narrow in the portion where the nascent RNA strand separates from the DNA template strand (Fig. 2c), measuring ∼16 Å in diameter. The backbone of the emerging RNA strand is directed out of the exit tunnel by the contacting residues R1020 and L315 of RPA2 and RPA1 respectively (Fig. 2c). Subsequently, the RNA exit tunnel widens up to ∼30 Å in diameter, conducive to the accommodation of the double-stranded RNA (Fig. 2d, e). The funnel formed by RPA1, RPA2 and the tWH domain of RPA49 is highly positively charged (Fig. 2e); its size and surface charge allow the formation of the RNA helix^39^. Indeed, the tunnel in human Pol I is wider, and the positively charged patch is larger compared to mammalian Pol II and Pol III (Fig. 2e). Also unique to Pol I is the position of the RPA49 tWH domain, which further extends the RNA exit tunnel with its positively charged surface (Fig. 2e).

**Fig. 2.**
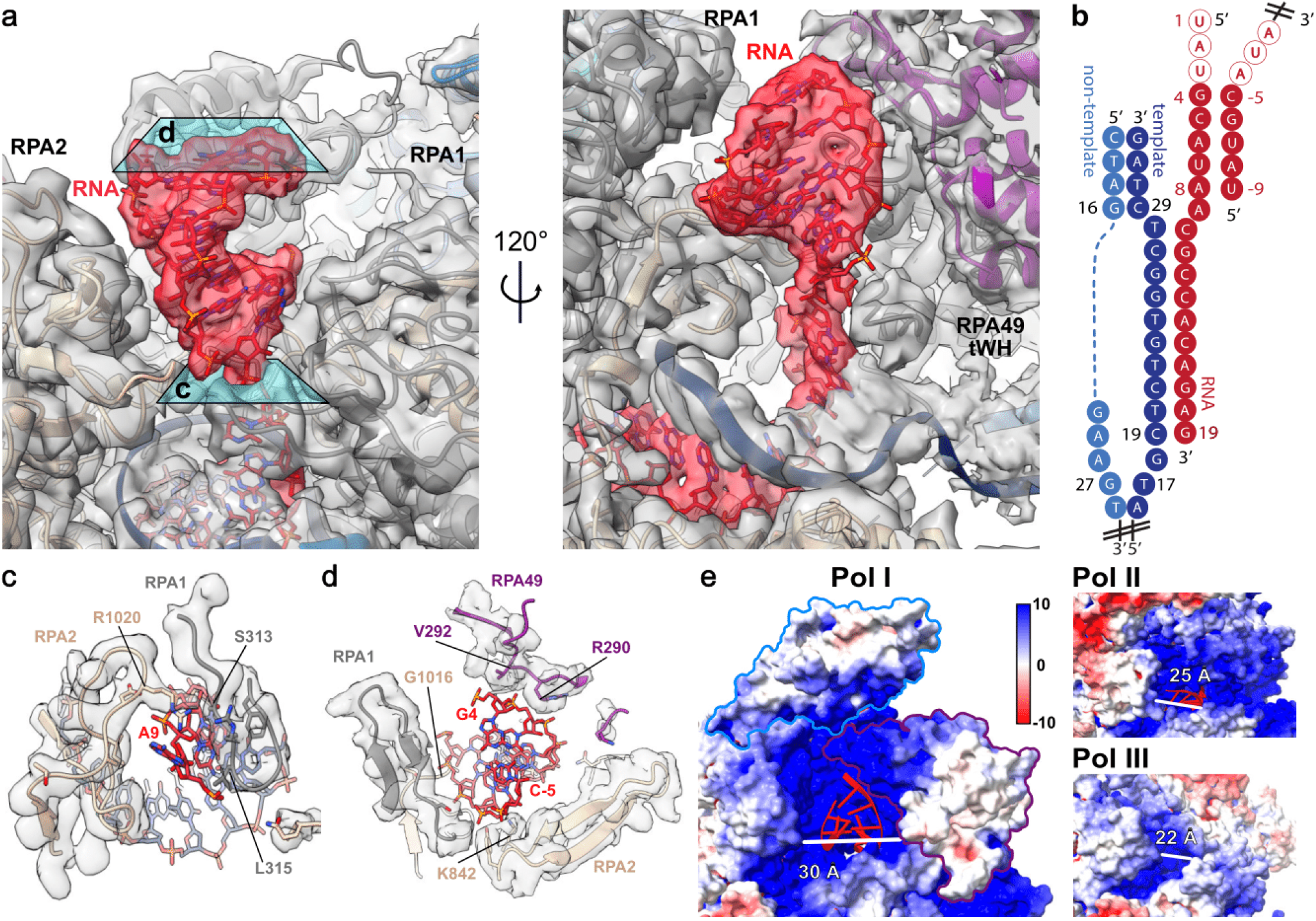
: RNA exit tunnel is able to accommodate a double-stranded RNA helix. **a** Close up view of the RNA exit tunnel. Cryo-EM density from Map B1 for the RNA (red) and the surrounding (grey) is shown. tWH domain of RPA49 is hidden in the left panel and a section of RPA2 subunit obscuring the RNA exit tunnel is hidden in the right panel. **b** Schematic representation of the modelled nucleic acid scaffold. Second RNA primer (right) base pairs to the primer annealed to the DNA scaffold forming a double-stranded helix due to the sequence complementarity. The empty circles correspond to the RNA bases not visible in the cryo-EM density. **c,d** Cross-section through the RNA exit tunnel at as indicated in the panel (**a**). Cryo-EM density corresponding to the protein components forming the tunnel is shown in grey. Residues within 5 Å of the RNA are shown in stick representation. Residues making contacts with the RNA are annotated. **e** Electrostatic charge distribution on the surface of the human Pol I EC (left panel), bovine Pol II (PDB: 5FLM^16^) (right top panel) and human Pol III (PDB:7AE1^17^) (right bottom panel). RNA in the exit tunnel is shown as cartoon (red). On the left panel, outlines of the RPA43 subunit (blue) and tWH of the RPA49 (purple) are shown. The tunnel width was measured using ChimeraX from backbone to backbone using: for Pol I residues RPA49 K362 to RPA1 S508, for Pol II residues RPB1 K434 to RPB2 Q838 and for Pol III residues RPC1 Y434 to RPC2 A798.

The specialized adaptations of human Pol I in the RNA exit tunnel may facilitate nascent RNA folding and thus positively influence the transcription rate^40^. Stable RNA structures correlate with high transcription rate since they prevent the transcript’s re-entry into the active site, which is required for backtracking. While the study by Turowski *et al.*^40^ used mainly yeast Pol I, the positive effect of RNA folding on elongation rates seems to be conserved among all RNA polymerases. Nevertheless, our results show that the human Pol I might possess a unique propensity to stabilize the RNA structures in the exit tunnel compared to human Pol III, where no similar RNA structure could be observed despite the use of the same RNA primer^17^. High transcription rates might be especially important for the transcription of the human rDNA repeat due to its transcribed length of 13 kb in humans compared to 6.9 kb in *S. cerevisiae*^1^. rRNA folds co-transcriptionally and undergoes complex processing along the ribosome biogenesis pathway^41^ requiring specific RNA structures. The wide and positively charged RNA exit tunnel observed in human Pol I allows the formation of such RNA structures.

### Human Pol I stalk contains a single subunit

The stalk of yeast Pol I consists of two subunits (Fig. 3a): the larger subunit A43, homologous to Rpb7 in Pol II and C25 in Pol III; and the smaller subunit A14, homologous to Rpb4 in Pol II and C17 in Pol III^18,19^. Nonetheless, sequence homology searches did not identify any homologous proteins to the yeast subunit A14 in humans. The cryo-EM structure of human Pol I reveals that the stalk solely contains RPA43 (Fig. 3b), whereas in the stalk of yeast Pol I, clear cryo-EM density corresponding to two helices of the subunit A14 is additionally visible (Fig. 3a, b). Instead in the human RPA43 subunit, the ordered N-terminal portion (residues 28-44) forms a helix which points away from the polymerase core, occupying the same space as the helices of yeast A14 in the stalk (Fig. 3c, d). The corresponding region in yeast A43 instead points in the opposite direction and is bound to the Pol I core by a 36-residue extension of subunit A135, which is absent in the corresponding human subunit RPA1 (Fig. 3c, d). Consequently, the human Pol I stalk is less tightly bound to the core compared to the yeast Pol I stalk. In addition, the lack of the structured helical extension in the N-terminal portion of subunit RPABC2 and the presence of more flexible loop (residues 527-533) in subunit RPA1 (Fig. 3e, f) permit tilting of the stalk towards the tWH domain of RPA49 (Fig. 3g, h).

**Fig. 3:**
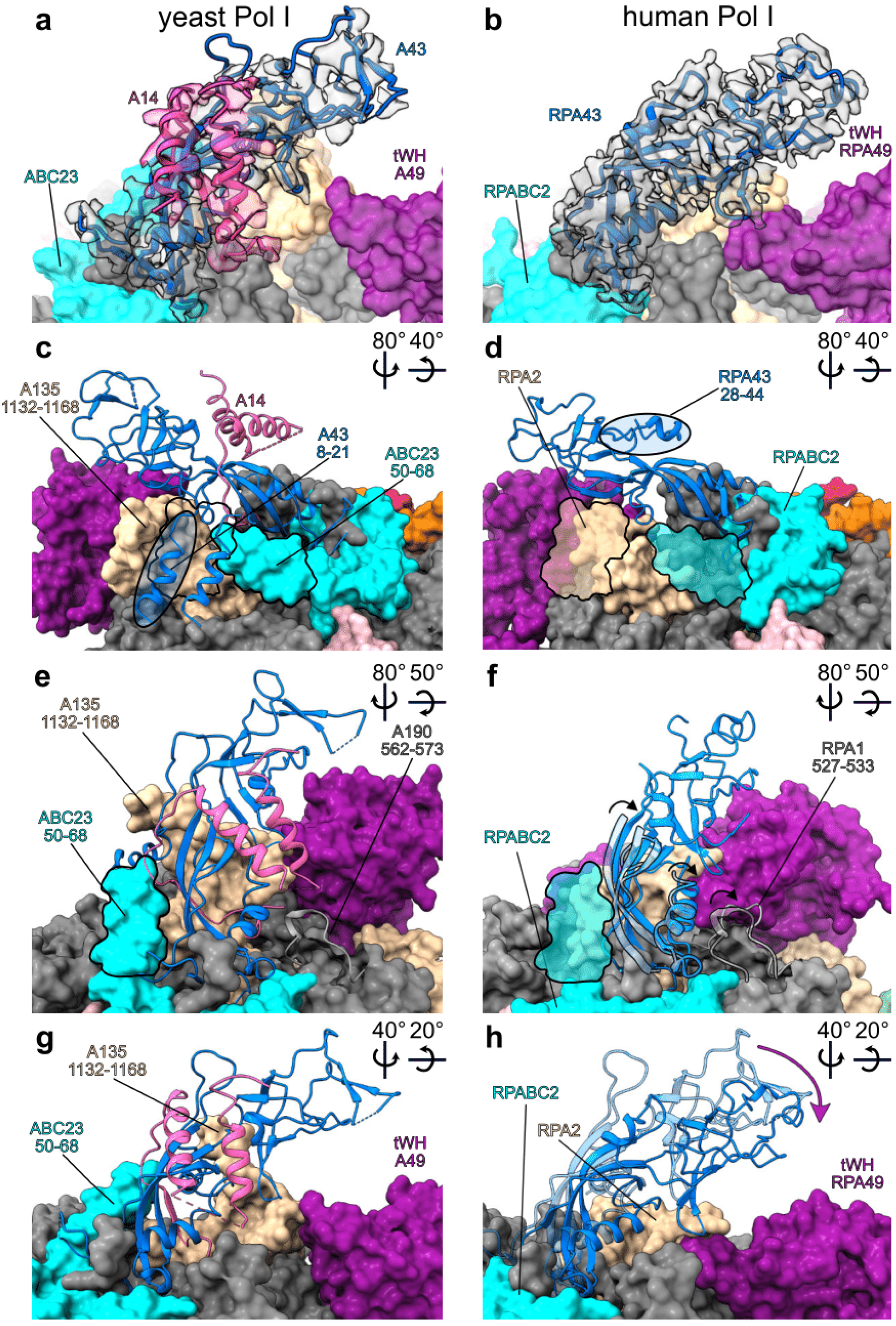
Human Pol I stalk is made out of one subunit only. **a, c, e, g** Yeast Pol I stalk (PDB: 5M64^28^ compared to **b, d, f, g** human Pol I stalk. Stalk subunits A43 (yeast) and RPA43 (human) (blue) and yeast A14 (pink) are shown in the cartoon representation, while the rest of the Pol I is shown in the surface representation colored by subunit as in Fig. 1. The rotations of the views between the panels are indicated with arrows in the top right corner of the panels. **a, b** Cryo-EM density corresponding to A43/RPA43 (grey) and A14 (pink) is shown in a transparent representation. **c, d** Structurally homologous N-terminal portions of the A43 and RPA43 subunits are marked with the blue transparent oval. **e, f** Structured in (**e**) yeast A190 loop (562-573) in cartoon representation is overlaid on (**f**) the human structure as a transparent cartoon. **f** In humans, the RPA1 loop (527-533) (cartoon representation, solid) is less structured and folds away from the stalk (bottom black arrow). Structural elements within the lower part of the stalk in humans tilt in the same direction (black arrows) when compared to yeast (overlaid transparent cartoon of the lower part of the yeast stalk). **c-f** Outlines of the yeast specific extensions in A135 and ACB23 marked in (**c** and **e**) are overlaid onto the human structure in (**d** and **f**) respectively. **g, h** In the human structure, the stalk leans (purple arrow) towards the RPA49 tWH domain, compared to the yeast structure, which is overlaid as a transparent cartoon.

The absence of an A14 homolog in humans prompted us to conduct an extended phylogenetic analysis (Extended Data Fig. 2, Supplementary Table. 2). We found that while the presence of the second, smaller stalk subunit is conserved throughout the eukaryotic tree of life for Pol II, in the case of Pol I it is specifically found only in some species from a division of Fungi, Ascomycota (Extended Data Fig. 2a). Thus, based on our structures and phylogenetic analysis, we show that Pol I in general contains 13 subunits, except in the few species of fungi. Consequently, previous studies using *S. cerevisiae*^19,20^ or *S. pombe*^42^ have focused on outliers within the tree of life. We then investigated the phylogenomic conservation of the yeast extensions of subunits A135 (1135-1168) and ABC23 (50-68) which contact the stalk (Fig. 3e). Since the subunits A135 and ABC23 are highly conserved across the tree of life, we created sequence alignments of the full-length proteins (Supplementary Files 1 and 2) and checked for the presence of insertions (Extended Data Fig. 2a). We found that the insertion in the A135 (1135-1168) appeared within Fungi (Extended Data Fig. 2a, b), which correlates with the presence of the two-subunit stalk in some species. The N-terminus of the ABC23 subunit is less conserved (Extended Data Fig. 2c, d) and thus its relation to the stalk subunits requires further investigation. The absence of an A14 homolog in human Pol I likely renders the stalk more flexible, which is consistent with our cryo-EM data and could play a role in the association of transcription factors such as RRN3 via conformational selection.

### Structured extensions bind the heterodimer to the core

The human Pol I heterodimer consists of subunits RPA34 and RPA49 and is structurally and functionally related to TFIIF/TFIIE in the Pol II system^29,30,43^. The overall architecture of the heterodimer consisting of a dimerization module and long extensions that bind to the core is conserved across species^19,36,37^. Yet, sequence identity of this subcomplex between yeast and human is only 20% and 25% for RPA49 and RPA34 respectively, which is the lowest among Pol I subunits (Fig. 1b). Subunit RPA34 in humans is over twice as large as in yeast (Extended Data Fig. 3a) owing to a large, disordered C-terminal tail. While the majority of this C-terminal extension is disordered in both yeast and humans, the first ∼40 amino acids (residues 120-161 in humans) can be assigned to a density running along the core of Pol I, and docking into a cleft within subunit RPAC1 (Extended Data Fig. 3c). While the general path of the extension is conserved across humans and yeast, its sequence is highly divergent (Extended Data Fig. 3b, c)^19,20^. In human RPA34, the extension is proline-rich and adopts a rigid conformation comprising of several kinks, which are contacted by core residues (Extended Data Fig. 3c close-up panels). Restricted conformational flexibility of proline-rich elements creates on one side a perfect hydrophobic stretch, while the opposite side harbors many sites accessible for hydrogen bonding, which together serve as a favorable platform for protein-protein interactions^44^. Instead, in yeast A34.5, the extension contains more charged residues (Extended Data Fig. 3b), which can contact core residues and form hydrogen bonds. Taken together, the different sequences accomplish the same function of tightly anchoring the heterodimer to the Pol I core.

Other possibly diverse roles of the disordered C-terminus of RPA34 are not fully understood. The C-terminal extension of yeast A34 harbors a nucleolar localization signal^45^. It has also been shown that human RPA34 can diffuse out of the nucleolus upon starvation^46^, which might be modulated by post-translational modifications within the C-terminal extension, for example, through acetylation^47^. The N-terminal end of RPA34 has been shown to interact with SL1, while the C-terminal extension appears to affect the rate of rRNA transcription and was also suggested to bind SL1 based on *in vivo* studies ^46^. RPA34 was also proposed to interact with UBF, possibly through multiple binding sites throughout the C-terminal extension^48^. In the human Pol I EC structure, only 150 residues of RPA34 are ordered with clear cryo-EM density, hence the remainder of the C-terminal extension may possibly become ordered upon binding to other Pol I factors.

Subunit RPA49 is a hybrid between TFIIF and TFIIE due to its N-terminal dimerization and C-terminal tWH domain, respectively (Extended Data Fig. 4a)^36,37^. N- and C-terminal ends of RPA49 are located on opposite sides of Pol I, with the N-terminal dimerization domain anchored to the RPA2 subunit and the C-terminal tWH domain binding the clamp close to the stalk and the RNA exit tunnel (Extended Data Fig. 4b). The overall fold and position of the human RPA49 tWH domain is similar to the one found in yeast A49 (Extended Data Fig. 4c). The tWH domain has DNA binding capability and may change its position to contact upstream DNA as seen in yeast Pol I PIC^30^. The linker following the dimerization domain and positioning the tWH domain runs along the upper clamp and is tightly anchored to a knob formed by the two coiled-coil helices in the RPA1 clamp core (Extended Data Fig. 4d left panel). The partially disordered loop (residues 345-383) connecting these two helices changes its conformation upon RPA49 linker binding (Extended Data Fig. 4d). The RPA49 linker further crosses the DNA binding cleft in close proximity to the downstream double-stranded DNA (Extended Data Fig. 4e, left panel). The RPA49 linker harbors a helix-turn-helix (HTH) motif (Extended Data Fig. 4e right panel) which is structurally and functionally conserved between yeast and humans, despite only partial sequence conservation (Extended Data Fig. 4f). It is predicted to bind DNA and was shown to be required for cell proliferation^49^. While mutational studies *in vitro* show that both helix 1 and helix 2 have DNA binding capability^49^, in our structure only helix 2 is poised for interactions with the DNA whereas helix 1 is bound to the RPA2 lobe (Extended Data Fig. 4e, right panel).

### Structure of human Pol I bound to RRN3 and in the Open Complex conformation

To further investigate the structure and function of human Pol I, we obtained the structure of Pol I bound to the initiation factor RRN3^50^. Human RRN3 interacts with SL1 and is thought to prime Pol I for transcription initiation^51^. We incubated purified native Pol I with recombinant human RRN3 in equimolar ratios in the presence of the previously used DNA scaffold devoid of RNA. After extensive classification of the acquired dataset (Extended Data Fig. 5), we obtained two cryo-EM maps corresponding to Pol I bound to RRN3 (Pol I-RRN3) and Pol I bound to the open DNA template termed Open Complex (Pol I OC) (Fig. 4a, b). In the Pol I-RRN3 complex, clear cryo-EM density corresponding to RRN3 is visible in the expected location close to the stalk (Fig. 4). Due to the flexibility of the upper clamp, focused refinement was used to improve the map quality in this region (Extended Data Fig. 5, Table. 1). Despite the lower local resolution, this allowed placing a human RRN3 homology model into the corresponding density with a good fit for the, mostly helical, secondary structure elements.

**Fig. 4:**
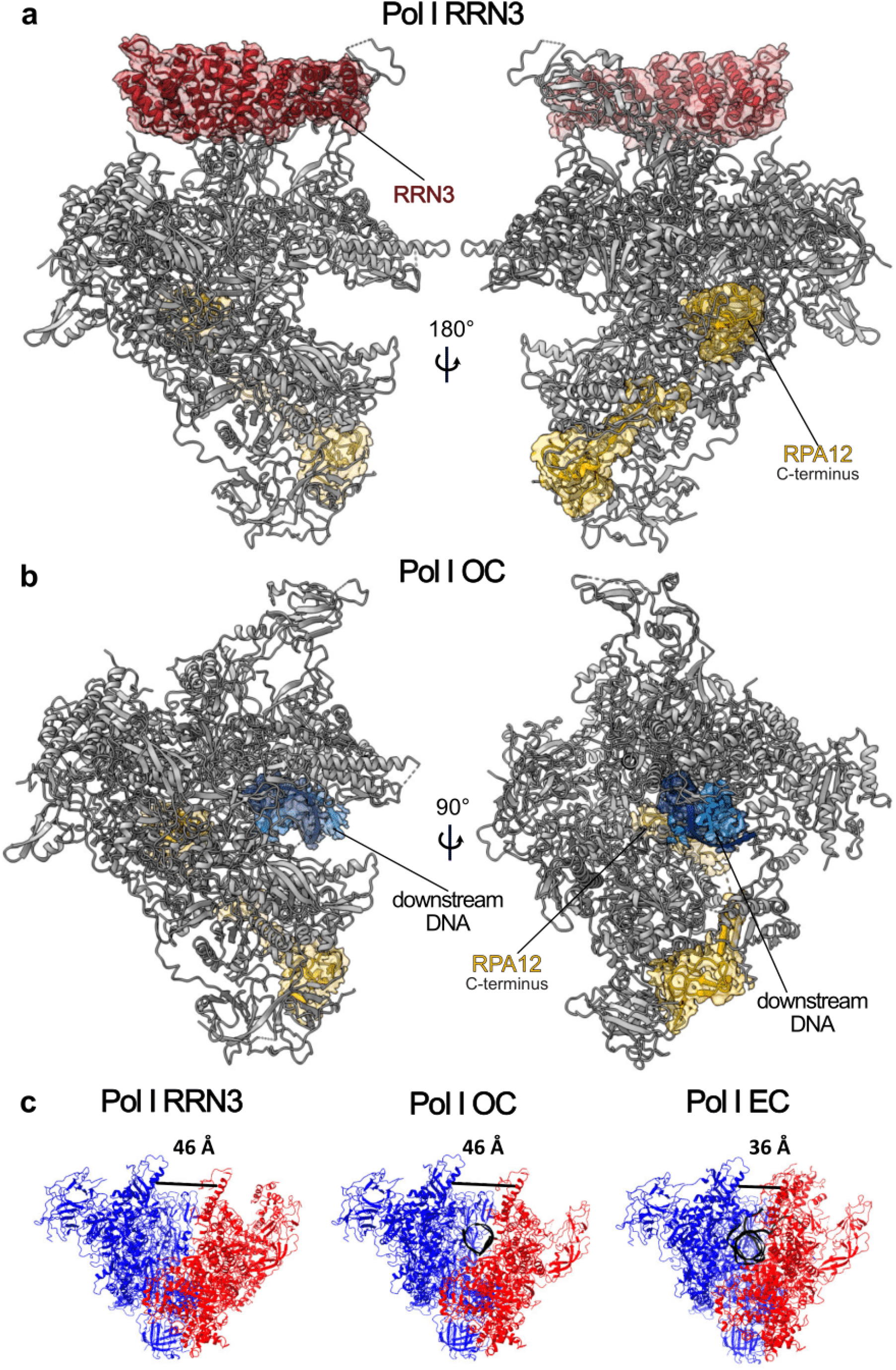
Structure of the Pol I-RRN3 and Pol I OC. **a, b** Pol I-RRN3 (**a**) and Pol I OC (**b**) are shown in cartoon representation. Cryo-EM density for RPA12 and DNA (in the Pol I OC) or newly built subunits (RRN3 in the Pol I-RRN3) is shown as transparent surface. **c** Differences in the cleft width between the three human Pol I structures. The two modules which move with respect to each other are colored in red (upper clamp and stalk) and blue (core and heterodimer). Nucleic acids are colored in black. The distance was measured from backbone to backbone of RPA1 L341 (clamp core) to RPA2 I396 (protrusion) and is shown with a black line.

The Pol I OC map has a slightly lower resolution (Table. 1), notwithstanding, we could confidently build the downstream portion of the DNA scaffold (Fig. 4b). In both structures, Pol I-RRN3 and Pol I OC, the RPA49 tWH domain is disordered (Fig. 4a, b), hence supporting the notion that it plays a role in clamp closing during transcription elongation^30,31,52^. Insertion of the C-terminal domain of the RPA12 subunit into the active site, observed in the Pol I-RRN3 and Pol I OC structures (Fig. 4a, b), might also be associated with the open clamp conformation. Closing of the clamp by up to 10 Å is associated with tight binding of the nucleic acids and elongation state, while in the transcriptionally inactive states represented by Pol I-RRN3 and Pol I OC, the clamp is more widely open and flexible (Fig. 4c). In yeast, a similar closing of the clamp has been observed, though the clamp closes by 7 Å between Pol I OC and EC states^28^ and by 6 Å between apo Pol I-dimer and Pol I OC states ^19,28^. So far, the high flexibility of the clamp hindered structure determination of human apo Pol I, and the dimerization of human Pol I has not been observed. At present, it therefore remains unknown if the Pol I clamp can open even further.

### Inactive state of Pol I

The opening of the clamp and insertion of the RPA12 C-terminal domain (corresponding to yeast A12 C-terminal domain) has been associated with the inactive state of Pol I^28^. The C-terminal domain of RPA12 is homologous to TFIIS in the Pol II system and possesses RNA cleavage activity^13,53^. Ordering of the C-terminal domain of A12 has previously been observed in the apo and OC conformations of yeast Pol I^19,20,28,33^, while it remains disordered in the EC state ^28,33^. In the human Pol I OC, the nucleic acid scaffold is bound at a distance from the active site in a non-productive conformation (Extended Data Fig. 6a, left panel). We could trace the double-stranded downstream DNA, but not the unwound portion of the DNA, indicating that it is not stably associated with the active site. In the Pol I OC, no density could be unambiguously assigned to the catalytic Mg^2+^ and the density for the catalytic aspartates is also less well defined compared to the Pol I EC (Extended Data Fig. 6a, b top middle panels). In line with the inactivation during cleft expansion mechanism suggested by Engel *et al.*^20^, one of the catalytic residues, D590 from RPA1, might have flipped out in comparison to the Pol I EC, which could contribute to the weaker cryo-EM density. The functional elements of the active site such as the bridge helix and the trigger loop are disordered, whereas these elements are fully ordered in the Pol I EC (Extended Data Fig. 6a, b right panels). The unfolding of the trigger loop in the Pol I OC is coupled to the insertion of the C-terminal domain of RPA12 subunit into the active site. The fully extended trigger loop as seen in the Pol I EC (Extended Data Fig. 6b, bottom right panel) would sterically clash with the C-terminal domain of RPA12 (Extended Data Fig. 6a, bottom right panel). The insertion of the RPA12 C-terminal domain also triggers a switch of the gating tyrosine (Y687 of RPA2). In the Pol I EC, the gating tyrosine occludes the backtracking funnel, and in Pol I OC, it flips forward to avoid the steric clash with residue D106 from RPA12 (Extended Data Fig. 6a, b middle bottom panels). Flipping of the gating tyrosine was first observed for yeast Pol II bound to TFIIS. However, in the model proposed by Cheung and Cramer^54^, the gating tyrosine flips to prevent its interaction with the backtracked RNA in the reactivation intermediate, thereby enabling TFIIS to fully insert into the funnel. Given that no RNA is present in the Pol I OC sample, we conclude that, at least for human Pol I, the insertion of RPA12, and not the backtracked RNA, induces flipping of the gating tyrosine.

Subunit RPA1 has a large insertion within its jaw (Extended Data Fig. 7a) which is fully disordered in the Pol I EC structure. In the yeast Pol I crystal structures, this insertion harbors the ‘DNA-mimicking loop’ or ‘expander’ that overlaps with the DNA backbone in the DNA-binding cleft^19,20^. In the Pol I-RRN3 conformation, we also observe weak cryo-EM density lining the cleft (Extended Data Fig. 7b) which would match the position of the DNA-mimicking loop in the yeast Pol I crystal structures. Like in yeast Pol I, when superimposed onto the Pol I EC, this extra density would clash with the DNA backbone (Extended Data Fig. 7c) suggesting that this density corresponds indeed to the DNA-mimicking loop and that this mode of regulation is conserved between yeast and humans. While there is little sequence identity between yeast and human Pol I in the DNA-mimicking loop region (Extended Data Fig. 7e), in both cases the insertion is negatively charged, suggesting a similar mode of function. Although the density was not of sufficient quality to build an atomic model of the complete human Pol I DNA-mimicking loop, we could unambiguously assign the register for the first 10 residues of the DNA-mimicking loop (1365-1375). When we superimposed its density onto the Pol I EC, it also overlaps with the turn from the HTH motif within the RPA49 linker (Extended Data Fig. 7d). Insertion of the DNA-mimicking loop into the cleft could therefore also prevent positioning of the RPA49 linker that assists in closing the clamp in the transition to transcription elongation^28,52^.

### RRN3 binding to Pol I affects the stalk

Human RRN3 binds to Pol I in a position similar to the one observed in yeast^29,30,32^. It contacts the stalk subunit RPA43 and residues 468-542 (dock) of subunit RPA1 (Fig. 5a). Superimposition with the Pol I EC suggests that the RPA49 tWH domain would clash with RRN3 (Fig. 5a). It has been suggested that the RPA49 tWH domain may help dislocate RRN3 in the transition from initiation to elongation^30,37^. Yet, a structure containing both RRN3 and A49 tWH bound to the upper clamp is available from yeast ^30^. In this structure, a larger portion of the RRN3 C-terminus is ordered, compared to the human Pol I-RRN3 structure. In humans, RPA49 can interact with the C-terminus of RRN3 ^55^, which might also subsequently become ordered at a later stage of the transcription cycle.

**Fig. 5:**
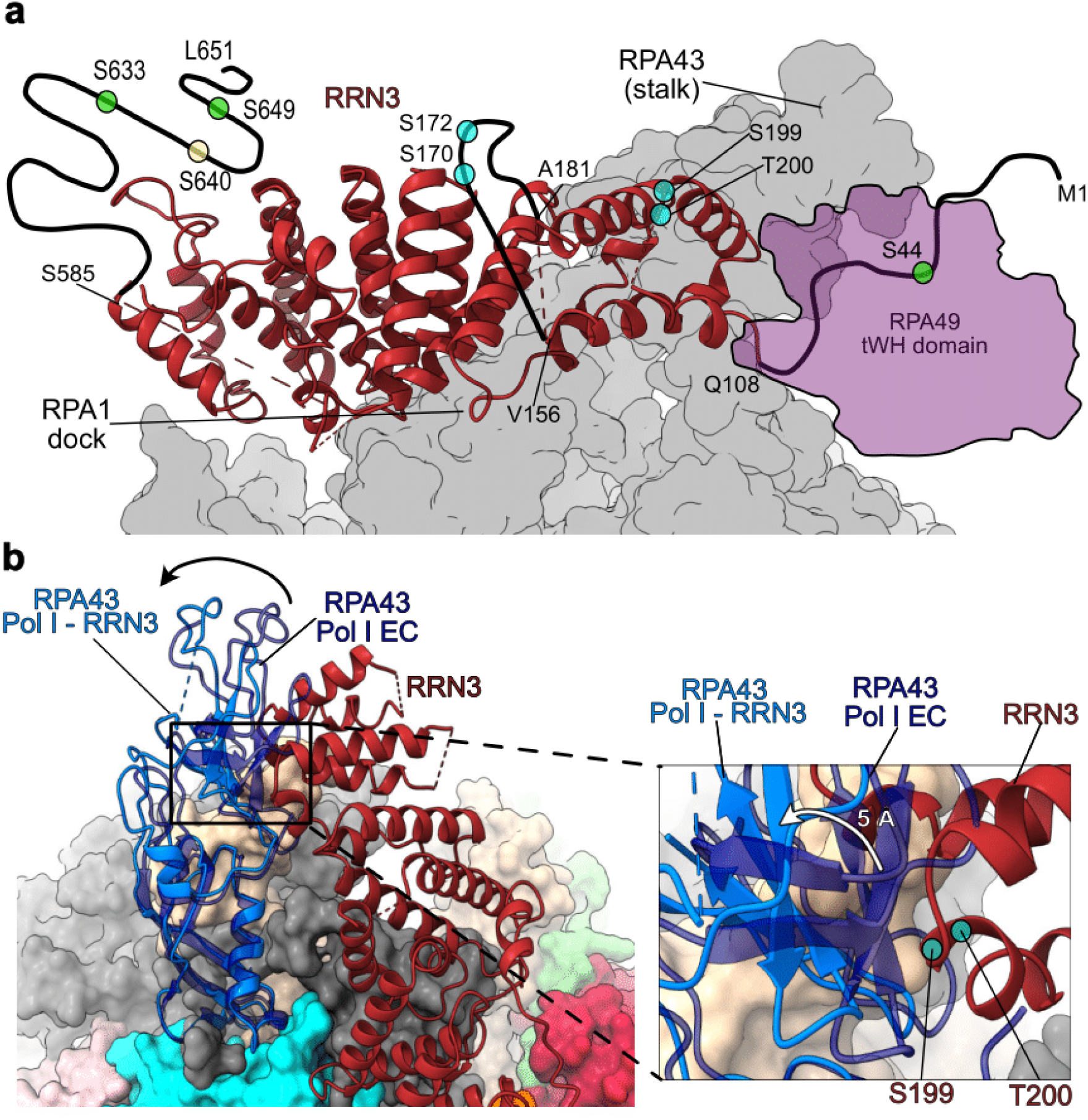
RRN3 binding to human Pol I. **a** RRN3 (maroon, cartoon representation) binds the Pol I (grey, surface representation) at stalk and RPA1 dock regions. Disordered parts of the human RRN3 which harbor phosphorylation sites are shown with black solid lines (not to scale). Phosphorylation sites are marked with circles: activating phosphorylation sites in green^26,27^, inactivating phosphorylation sites in cyan^25,26,56^ and phosphorylation site of unknown role in yellow^57^. The position of the RPA49 tWH domain from the Pol I EC is shown with a purple, transparent outline. **b** RPA43 subunit and RRN3 are shown in cartoon representation while the rest of the Pol I is shown in surface representation colored according to the subunit as in Fig. 1. Upper part of the RPA43 subunit from the Pol I-RRN3 structure (light blue) swings away (black arrow) from the RRN3, compared to the RPA43 subunit from the Pol I EC structure (dark blue, transparent). The lower part remains anchored to the core. The closest contact point between RPA43 and RRN3 (close-up view) is the location of the two residues (S199 and T200) which can carry an inactivating phosphorylation (cyan circle). RPA43 from the Pol I EC complex (dark blue transparent) would clash with the RRN3 and thus it swings by 5 Å away (white arrow) into the Pol I-RRN3 conformation.

In human Pol I, the main interaction site for RRN3 is the RPA43 subunit. The absence of a human subunit equivalent to yeast A14 introduces a hinge in the middle of the stalk-forming RPA43 subunit. This hinge allows the upper part of the stalk to swing away by ∼5Å upon RRN3 binding, while the bottom part of the stalk remains anchored to the Pol I core (Fig. 5b). Binding of RRN3 to human Pol I for promoting transcription initiation relies on the phosphorylation status of RRN3^25–27,56,57^. The majority of residues that can be phosporylated lie in disordered regions of RRN3 (Fig. 5a), which may become ordered upon interaction with SL1 or UBF. However, two residues (S199, T200) that were shown to be a target of inactivating phosphorylation lie in the interface with subunit RPA43 (Fig. 5b right panel). We speculate that phosphorylation at these two positions, adjacent to each other, may function as a phospho-switch and disrupt the interaction between RRN3 and RPA43 (Fig. 5b right panel). RRN3 also needs to be phosphorylated at several other sites to be able to bind to Pol I and stimulate specific transcription initiation^58^. They are key for regulation of Pol I activity in response to proliferation signals^27^, nutrient availability^26^ or stress^25^. In addition to phosphorylation events that directly affect the affinity between RRN3 and Pol I, some of these phosphorylations might also stimulate human Pol I activity by increasing the affinity of RRN3 to the accessory factors SL1 and UBF.

### Disease-associated mutations in Pol I

Disorders known as ribosomopathies are associated with Pol I transcription and its associated factors^7^. Several disease-causing mutations are found within Pol I itself. They cluster in subunits RPAC1 and RPAC2 shared between Pol I and Pol III, as well as around the active site of Pol I (Fig. 6a). Mutations within subunit RPA2 cause Teacher Collins Syndrome (TCS), which is a craniofacial developmental disorder^9^. Residue S682 contacts residues within the bridge helix (Fig. 6b), while residue R1003 lies within the hybrid binding region and interacts with the fork (Fig. 6c). Precise positioning of those active site elements throughout the transcription cycle is required for Pol I function and thus alterations in these regions could greatly impair Pol I activity. Mutations causing Acrofacial Dysostosis (AD), another disease with craniofacial abnormalities that arise during development, are found within the RPA1 subunit^8^. Residue E593 is found adjacent to the catalytic aspartate triad within the active site (Fig. 6d), and changes to this site could interfere with nucleotide addition. Residue V1299 is located away from the active site, within the jaw domain and yet it contacts the RPA12 linker (Fig. 6e). This interaction could be important for the correct insertion of the RPA12 C-terminal domain into the active site and thus influence Pol I inactivation or backtracking. A large number of TCS causing mutations are found in the RPAC2 subunit which is important for core stability (Fig. 6f)^59,60^. Since the subunit RPAC2 is shared between Pol I and Pol III, it is possible that mutations in it affect both Pol I and Pol III. Another TCS causing mutation, R279 found in subunit RPAC1, is also shared between Pol I and Pol III, but mostly affects Pol I. In human Pol III, the mutated residue R279 is highly exposed to solvent and does not make many contacts with other subunits (Fig. 6g, right panel)^17^. In human Pol I on the contrary, R279 binds to the RPA34 extension that buries this residue (Fig. 6g, left panel). Using immunofluorescence and affinity pull-downs it was shown that R279 does not affect assembly of Pol III, while it influences the localization of Pol I to the nucleolus^61^. Therefore, the impaired nucleolar localization or assembly of Pol I presumably results in the TCS-causing effect of this mutation. Subunits RPAC1 and RPAC2, as well as Pol III-specific subunits, also harbor many mutations causing Hypomyelinating Leukodystrophy (HLD)^12,17,61^. Some of those mutations cluster in the C-terminus of the RPAC1 subunit (residues 26-32)^12^, which is fully ordered in human Pol III, hence making it an important interaction site with other core subunits (Fig. 6h, right panel). However, the RPAC1 N-terminus (1-37) is disordered in human Pol I (Fig. 6h, left panel). Thus, HLD-associated mutations in this region cannot directly impair the activity of Pol I. In line with our observations, it was shown that the mutation of one of those residues (N32) as well as one residue within the core of RPAC1 (N74) impair the assembly of Pol III, but not Pol I^61^. Our structure suggests that HLD is likely to arise from perturbations to Pol III and not Pol I. On the contrary, TCS is more likely a result of Pol I malfunction. The high-resolution structure of human Pol I can help to explain the consequences of mutations on the molecular level and can direct further studies of disease-associated mutations. Together with the structure of human Pol III^17^ it also enables distinguishing effects of mutations in subunits that are shared between Pol I and Pol III.

**Fig. 6:**
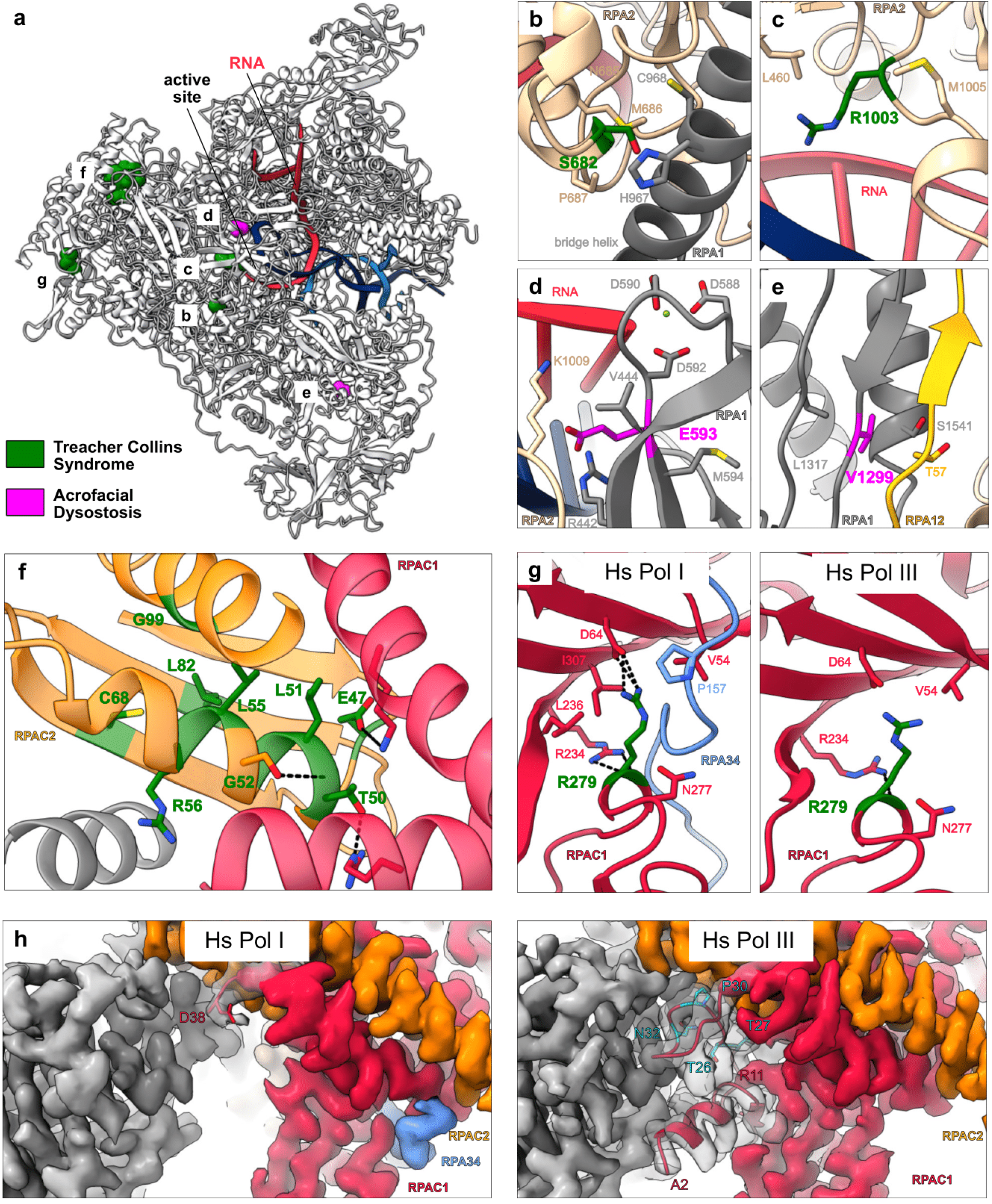
Disease associated mutations of Pol I. **a** Overview of the mutations affecting Pol I. The structure is shown in cartoon representation (grey) with nucleic acids marked in blue (DNA) and red (RNA). Disease-causing mutations for TCS (green) and AD (magenta) are shown in sphere representation. **a-g** Close-up views of the residues affected by the mutations (stick representation). Other Pol I residues that can make contacts with the mutated one are also shown in stick representation. Putative hydrogen bonds are shown with black, dashed lines. **d** Active site residues found in close proximity to the E593 residue are shown in stick representation. **f** Only residues affected by mutations and those that can form hydrogen bonds with them are shown for simplicity. **g** RPAC1 residue R279 is shown in human Pol I (left panel) and in human Pol III (PDB: 7AE1^17^) (right panel). **h** Cryo-EM density corresponding to the core (grey) of the human Pol I (left panel) and human Pol III (right panel). Subunit RPAC1 is shown in red, RPAC2 in orange and Pol I-specific RPA34 in blue. N-terminal residues are shown in cartoon representation together with their corresponding density in grey, transparent representation. The first visible residue in Pol I (D38) is shown in stick representation. Residues mutated in HLD found in the RPAC1 N-terminus (visible in Pol III) are shown in stick representation (cyan) (right panel).

## Conclusions

In this study, we present high-resolution structures of the human Pol I in its elongating state at 2.7 Å resolution, bound to RRN3 at 3.1 Å resolution and bound to an open DNA scaffold at 3.3 Å resolution. Despite the conservation of the general architecture of Pol I throughout the tree of life, several features distinguish human Pol I from yeast Pol I. First, we show that human Pol I can accommodate double-stranded RNA in the exit tunnel, what might increase the elongation speed^40^. In addition, the large funnel at the end of the RNA exit tunnel might assist the co-transcriptional folding of rRNA.

The exact subunit composition of human Pol I was previously unknown, as the human homolog of yeast stalk subunit A14 had not been identified. We now show that the human Pol I stalk is formed by a single subunit. In fact, a single-subunit stalk in Pol I is conserved throughout the tree of life with the exception of certain fungi including yeast. Thus, we suggest that Pol I should be generally referred to as a 13-subunit complex. The advantages of having a smaller and more flexible stalk that we observe in human Pol I needs to be further investigated. Currently, we could show that it allows a change in conformation upon binding of accessory factors such as RRN3, what may play a role in the transcription cycle. Therefore, the human Pol I structure provides insights into basic mechanisms of Pol I transcription and association with transcription factors. Production of rRNA is a fundamental cellular process, and thus high conservation of many features of Pol I is expected across species. Stable association of the heterodimer related to the TFIIE-TFIIF transcription factors to the Pol I core is based on the same principle of long extension in both yeast and humans. The structures of Pol I-RRN3 and Pol I OC represent inactive states of human Pol I with RPA12 being inserted into the funnel, an unfolded bridge helix and trigger loop, and a conserved (yet weakly bound) DNA-mimicking loop. Thus, critical elements that can regulate Pol I activity and render it inactive are also conserved between yeast and humans.

The structure of human Pol I also sheds light on the molecular basis of mutations that cause developmental disorders such as TCS, AD or HLD. We could map Pol I-specific mutations in its two largest subunits at areas close to the active site or in the RPA12 binding region. By comparison to the human Pol III structure^17^, we can rationalize the observation made by Thiffault *et al.*^61^ suggesting that HLD and TLD might arise from perturbations to Pol III and Pol I, respectively; even though the causative mutations lie in subunits shared between both polymerases. Therefore, the human Pol I structure better explains the effect of disease-causing mutations and lays ground for further investigations of disease mechanisms.

So far, detailed structural insights into human Pol I have been lacking. Our study highlights the considerable differences between yeast and human Pol I. Human Pol I and its general transcription factors are well-established targets of anti-cancer drugs^3^. We are convinced that this high-resolution cryo-EM structure of human Pol I will aid the design of additional, highly specific human Pol I-specific inhibitors. Additionally, access to the endogenously sourced pure human Pol I opens avenues for high-throughput drug screening as well as structural studies and lead optimization of small-molecule inhibitors, further supporting efforts to combat cancer.

## Acknowledgments

We thank F. Weis and W.J.H. Hagen (EMBL Cryo-Electron Microscopy Service Platform) for the EM support, P. Haberkant and M. Rettel (EMBL Proteomics Core Facility) for carrying out MS analysis, F. Dossin for help with CRISPR-Cas9, T. Hoffmann and J. Pecar for setting-up and maintaining the high-performance computing environment, A. Bateman (EMBL-EBI) for help improving Pfam families, and all current members of the Müller lab and S. Eustermann for discussions. A.D.M., M.G. and A.L. acknowledge support by the EMBL International PhD program.

## Author Contribution

C.W.M. initiated and supervised the project. A.D.M. designed the experiments, generated the CRISPR-Cas9 engineered HEK293T cell line, purified human Pol I, carried out EM grid preparation, data collection and processing, built the atomic models and interpreted the structures. M.G. advised on EM data processing, built the atomic models and interpreted the structures. H.G. performed cell culture work, advised and assisted with CRISPR-Cas9 mediated gene-editing and protein purification. F.B. did *in vitro* transcription assays. B.M. expressed and purified human RRN3. A.L. performed the phylogenomic analysis. A.D.M., M.G. and C.W.M. wrote the manuscript with input from the other authors.

## Methods

### Endogenous tagging of RPAC1 with CRISPR-Cas9

The cell line with homozygously tagged RPAC1 generated in Girbig *et al.*^17^ was used for purification of the human Pol I. Briefly, HEK293T cells were transfected using polyethylenimine reagent in Opti-MEM I Reduced Serum Medium (Gibco) with a pSpCas9(BB)-2A-GFP (PX458) plasmid (Addgene plasmid #48138)^62^ containing a guide RNA (5’-ACTGAGCTTGGATGCTTCTG-3’) and a donor plasmid containing 700 bp homology arms and a RPAC1 tag synthesised by GenScript into pUC57-Mini plasmid. The C-terminal RPAC1 tag contained mCherry, Strep II, 6xHis and P2A peptide followed by a blasticidin resistance gene and a SV40 termination signal. Cells were selected 5 days after the transfection with 5 µg/mL blasticidin (Thermo Fisher Scientific). They were then seeded at low density and single cell colonies were picked and genotyped by PCR. A clone with correct tag integration was adapted to growth in suspension. For assessing the localisation of the tagged complex, adherent cells were seeded to 20% confluency 2 days before imaging and supplemented with 1 μg/mL live Hoechst dye. The medium was replaced with fresh one immediately before imaging. Cells were imaged using a confocal microscope Olympus FV3000 using an UPLSAPO 60X silicone objective. Hoechst staining was detected with 405 nm excitation laser and mCherry was excited with 561 nm laser. Images were processed using Fiji^63^.

### Purification of human Pol I

Cells with tagged RPAC1 were grown in suspension in Expi293 Expression Medium (Thermo Fisher Scientific) up to 7 x 10^6^ cells/mL. They were harvested by centrifugation followed by washing with PBS and the pellets were flash-frozen for storage. Approximately 50 g of cell pellets were resuspended by stirring in lysis buffer (25 mM HEPES pH 7.5, 150 mM (NH_4_)_2_SO_4_, 5 mM MgCl_2_, 5% glycerol, 20 mM imidazole, 0.5% Triton-X100, 2 mM β-mercaptoethanol in the presence of EDTA-free protease inhibitor cocktail (Roche) and benzonase (Sigma-Aldrich)). Cells were sonicated, centrifuged for 1 h at 235,000 g at 4°C and filtered. The cleared lysate was applied to a 5 mL HisTrap HP column (GE Healthcare) and washed with Ni wash buffer containing 25 mM HEPES pH 7.5, 150 mM (NH_4_)_2_SO_4_, 5 mM MgCl_2_, 5% glycerol, 20 mM imidazole and 2 mM β-mercaptoethanol. The complex was eluted with a 300 mM imidazole step. Eluted fractions were combined and incubated for 1 h at 4°C with Strep-Tactin agarose beads (IBA Lifesciences). Beads were applied to a gravity column, washed with buffer containing 25 mM HEPES pH 7.5, 200 mM CH_3_CO_2_K, 5 mM MgCl_2_, 5% glycerol, 2 mM β-mercaptoethanol and eluted in 3 fractions with a buffer containing 20 mM biotin. Obtained fractions were pooled and applied to a MonoQ column (GE Healthcare). A gradient from 200 mM to 2 M CH_3_CO_2_K was used for elution and Pol I eluted in two fractions at approximately 40 mS/cm and 50 mS/cm. The 1^st^ peak was used for further studies. Pol I was concentrated on 100K spin concentrator (Merck Millipore) up to 0.7-1 mg/mL and buffer exchanged into the EM buffer containing 15 mM HEPES, pH 7.5, 80 mM (NH_4_)_2_SO_4_, 5 mM MgCl_2_ and 10 mM DTT. Quality of the complex was assessed with SDS-PAGE stained with Coomassie blue and the identity of all subunits was confirmed with mass spectrometry. The obtained sample was directly used for grid preparation or flash-frozen and stored for *in vitro* assays.

### Purification of RRN3

N-terminal His-tagged human RRN3 was cloned into pETM11 plasmid (EMBL Protein Expression and Purification Core Facility). It was transformed into *E. coli* LOBSTR expression strain (Kerafast). Cells were grown overnight at 18°C in TB medium and expression was induced with 0.05 mM of IPTG at OD_260nm_ = 0.8-1.0. Pellets were harvested by centrifugation and resuspended in lysis buffer (50 mM Tris-HCl pH 7.5, 200 mM NaCl, 10% glycerol, 10 mM imidazole, 2 mM β-mercaptoethanol in presence of Dnase 1 (Roche), EDTA-free protease inhibitor coctail (Roche) and lysozyme (Sigma-Aldrich)). Cells were disrupted using Microfluidizer Processor M-110L (Microfluidics) followed by centrifugation. The supernatant was incubated for 1 hour with Ni-NTA agarose beads (Qiagen) at 4°C. The agarose beads were then first washed with wash buffer 1 (50 mM Tris-HCl pH 7.5, 1 M NaCl, 10% glycerol, 40 mM imidazole, 2 mM β-mercaptoethanol and 5 mM ATP), followed by wash buffer 2 (50 mM Tris-HCl pH 7.5, 200 mM NaCl, 10% glycerol, 10 mM imidazole, 2 mM β-mercaptoethanol) and eluted with the elution buffer (50 mM Tris-HCl pH7.5, 200 mM NaCl, 10% glycerol, 150 mM imidazole, 2mM DTT). Eluted RRN3 was dialyzed in buffer A (20 mM Tris-HCl pH 7.5, 200 mM NaCl, 2 mM DTT) and the N-terminal His tag was cleaved off by overnight incubation at 4°C with TEV protease (EMBL Protein Expression and Purification Core Facility). The complex was then incubated for 30 min with the Ni-NTA beads (Qiagen) at 4°C to capture the cleaved off tag and the protease. The flow through was collected and applied to a MonoQ column (GE Healthcare). The RRN3 was eluted with a gradient over 10 column volumes of buffer B (20 mM Tris-HCl pH 7.5, 1 M NaCl, 2 mM DTT). The fractions containing RRN3 were concentrated using a 5 kDA cut-off concentrator (Corning) and injected on to a Superdex 200 increase 10/300GL size exclusion column (GE Lifesciences) pre-equilibrated with the gel filtration buffer (25 mM Tris-HCl pH7.5, 150 mM NaCl, 2 mM DTT). The fractions containing pure human RRN3 were concentrated up to 10-15 mg/mL and flash-frozen in liquid nitrogen for storage.

### Nucleic acid scaffold preparation

Nucleic acid oligonucleotides (Sigma-Aldrich, HPLC-grade) used included: template DNA: 5’ GTACTGAATTAGACAATGCTCTGTGGCTCTAGTACCATGAGCG 3’; non-template DNA: 5’ CGCTCATGGTACTAGGCTTCGGAGAAGTTGTCTAATTCAGTAC 3’ and RNA: 5’ UAUGCAUAACGCCACAGAG 3’. Template and non-template DNA at 100 μM in H_2_O were mixed and heated up to 95°C for 2 min. They were immediately transferred to ice for 5 min incubation and supplemented with 2x hybridisation buffer (40 mM HEPES pH 7.5, 24 mM MgCl_2_, 200 mM NaCl, 20 mM DTT). RNA at 100 μM was heated on a 55°C heating block for 1 min and added to the DNA scaffold on ice. Resulting DNA-RNA mixture was brought to room temperature. For the Pol I OC complex preparation, steps with the RNA were omitted.

### *In vitro* transcription assay

The RNA primer was labelled at the 5’ terminus with [γ-^32^P]ATP using T4 PNK (New England Biolabs) and PAGE purified. It was assembled with the DNA scaffold as outlined above. For the RNA extension assay, 2 pmol of the radioactively labelled nucleic acid scaffold were incubated with 3 pmol of Pol I for 10 min at room temperature. Subsequently, 0.2 mM of NTPs in the buffer containing 20 mM HEPES pH 7.5, 60 mM (NH_4_)_2_SO_4_, 10 mM MgSO_4_ and 10 mM DTT were added to the reaction mixture before incubation for 45 min at 37°C. The reaction was stopped by addition of formamide. Samples were heated for 3 min at 95°C and loaded on a denaturing 17% gel (17% Acrylamide/Bis 19:1, 8M Urea, TBE 1%). The radioactive products were recorded using phosphor-imaging screens (Fujifilm) for capturing digital images.

### Phylogenomic analysis

Reference proteomes for selected species (listed in the Supplementary Table 2) were downloaded from the UniProt website (version 2021_02)^64^. The phylogenetic tree of eukaryotic species was obtained from the NCBI Taxonomy Database^65^. Homologs for proteins of interest in this study were retrieved with the HMMER tool (v3.2.1)^66^, using individual protein sequences and domain families from the Pfam database (version 34.0)^67^. The following UniProt and Pfam identifiers were used for protein homology searches: A14 (UniProt:P50106, Pfam:PF08203), A43 (UniProt:P46669, Pfam:PF17875), ABC23 (UniProt:P20435, Pfam:PF01192), A135 (UniProt:P22138, Pfam:PF00562) and RPB4 (UniProt:P20433, Pfam:PF03874). The Pfam family for A14 (Pfam:PF08203) has been improved to include the *S. pombe* homologous protein, previously missing from the model. Paralogs for homologs of RPB4 subunit have been assigned according to their similarity to the *S. cerevisiae* protein (UniProt:P20433). For the A43 subunit, a new HMM model was created with HMMER based on the multiple sequence alignment, generated using the MUSCLE tool (v3.8)^68^, of A43 proteins in six species: *H. sapiens, S. cerevisiae, D. melanogaster, A. thaliana, D. discoideum* and *S. pombe*. The Pfam family of the A43 OB domain (Pfam:PF17875) has been iterated to include missing domain annotations in higher Eukaryotes. Structural insertions in A135 and ABC23 were manually assigned based on a multiple sequence alignment of full protein homologs generated using MUSCLE^68^. The Interactive Tree of Life (iTOL) online tool^69^ was used to visualize and annotate the phylogenetic tree and create the final figure. Sequence consensus score and occupancy score across the ABC23 subunit consensus sequence in Extended Data Fig. 2 was generated using Jalview^70^. Multiple sequence alignments in Extended Data Fig. 3, 4 and 7 were obtained using Clustal Omega^71^.

### Cryo-EM sample preparation

Freshly purified Pol I was used to prepare cryo-EM grids. For the Pol I EC sample, annealed DNA-RNA nucleic acid scaffold in 1.5 molar excess was added to the Pol I at 0.7 mg/mL. For the Pol I-RRN3-OC sample, Pol I at 0.85 mg/mL was mixed with RRN3 (diluted in EM buffer) in equimolar ratio and incubated on ice for 20 min prior to the addition of the DNA OC scaffold in 1.5 molar excess. Both samples were incubated for 30 min at room temperature and then kept on ice until plunge freezing. Mesh 200, Cu R2/1 grids (Quantifoil) were plasma cleaned using a NanoClean plasma cleaner (Fischione Instruments, Model 1070) with 75% - 25% argon-oxygen mixture for 30 s. Grids were then plunge-frozen in liquid ethane using Vitrobot Mark IV (Thermo Fisher Scientific) set to 100% humidity and 15 °C. 2.5 μL of sample was applied to the grid and it was blotted using the following parameters: blot force 3, blot time 0 s, wait time 0 s.

### Data collection (Pol I EC)

Pol I EC dataset was collected on a Titan Krios TEM operated at 300 keV (Thermo Fisher Scientific) equipped with a K3 direct detector (Gatan) and a Quantum energy filter (Gatan) using SerialEM^72^. Magnification of 105,000x corresponding to a pixel size of 0.822 Å/pixel was used. We recorded 10,053 movies in counting mode with 1.77 e^-^/Å^2^/frame over 38 frames with a defocus range of 1.0-2.5 μm.

### Data collection (Pol I-RRN3 and Pol I OC)

Grids with Pol I-RRN3-OC sample were pre-screened on FEI Talos Arctica microscope equipped with a Falcon III detector. A small dataset of 967 stacks was recorded with SerialEM^72^ at a magnification of 92,000x corresponding to a pixel size of 1.566 Å/pixel. Exposure of 3.62 e^-^ /Å^2^/frame over 12 frames was used with a defocus range of 1.0-2.5 μm.

High resolution dataset was acquired as for the Pol I EC (Titan Krios TEM operated at 300 keV with magnification of 105,000x corresponding to a pixel size of 0.822 Å/pixel), but 14,224 movies with 40 frames and exposure of 1.03 e^-^/Å^2^/frame were collected. Defocus range of 0.75-2.25 μm was used.

### Data processing (Pol I EC)

The overall processing pipeline for the Pol I EC is outlined in Extended Data Fig. 1. WARP 1.0.7W^73^ was used for the initial pre-processing of micrographs: frame alignment, contrast transfer function (CTF) estimation and dose weighting. 2,171,736 particles were picked with the network BoxNet2Mask_20180918 and extracted with a box size of 280 pixels. Particles were imported into CryoSPARC^74^ and subjected to 2D classification. We selected 20 out of 200 classes with clear 2D class averages showing secondary structure features. Selected 942,771 particles were subjected to *ab initio* classification. One out of two produced models resembled the yeast Pol I and it was refined using Homogeneous Refinement (Legacy). Obtained map reached the overall resolution of 3.4 Å and was further used as a reference. Further processing was performed with RELION 3.1^75^.

All micrographs were motion-corrected using RELION’s own algorithm according to MotionCor2^76^ and CTF-corrected using Gctf^77^. Particles selected by WARP were re-extracted with a box size of 70 pixels (binned by factor of 4). The reference obtained from cryoSPARC was appropriately re-sized using EMAN2 package^78^ and low-pass filtered to 40 Å in RELION. The reference was used in a global 3D classification using T-parameter of 20. T-parameter = 20 was used in all 3D classification steps for both datasets. 3 out of 8 classes which contained a complete Pol I map were selected and 769,742 particles were re-extracted into a box size of 160 pixels (binned by factor of 2) and 280 pixels (unbinned). Unbinned particles were CTF-refined and subjected to Bayesian polishing^79^ giving a map that resolved up to 2.8 Å. While the map reached high resolution, it had rather poor density in the upper clamp and stalk region. Into the obtained map we fitted a human Pol I homology model based on the yeast Pol I crystal structure (PDB: 4C3I)^19^ and using UCSF Chimera^80^ we created a general soft mask using the molmap command. Next, we performed masked 3D classification using the general soft mask. One of the two obtained classes with 198,822 particles had more high-resolution features throughout the map and it was thus CTF-refined, polished and CTF-refined a second time. The obtained map denoted as Map A has an overall resolution of 2.7 Å.

To improve the density in the stalk region, particles selected after the 1^st^ global 3D classification (binned by factor of 2), were subjected to masked 3D classification with a soft mask in the stalk region. One of the two classes with 222,815 particles showed improved density in the region of interest and resolved up to 3.7 Å. Additional fuzzy density close to the stalk could also be observed at higher threshold levels. Guided by yeast elongating Pol I structure (PDB:5M64)^28^, we fitted a homology model for the RPA49 tWH domain. A soft mask covering the stalk and the tWH of RPA49 was used for another masked 3D classification. A class containing 37,180 particles showed improved density in the region and had an overall resolution of 3.9 Å. The same strategy of CTF-refinement and particle polishing was employed and the resolution improved up to 3.0 Å, giving Map B. Using the fitted homology model of human Pol I, two soft masks were created using Chimera: 1^st^ covered the upper clamp, stalk, tWH of RPA49 and the nucleic acid scaffold and 2^nd^ contained the core of the Pol I and the heterodimer. Those masks were used for multi-body refinement in RELION^81^ which yielded partial maps B1 and B2. The resolution of all maps was obtained with RELION’s post-processing tool based on the gold-standard Fourier shell correlation (FSC) using the 0.143 cut-off criterion^82^. The local resolution range was estimated using RELION’s local resolution tool. Map C is a composite map obtained using the “combine_focused_maps” command implemented in the Phenix 1.18^83^ supplied with the Map A, B and B1. Map A was sharpened with the LocalDeblur tool^84^ from Scipion^85^, while maps B, B1, B2 and C were sharpened with the Autosharpen tool from Phenix^86^.

### Data processing (Pol I-RRN3 and Pol I OC)

The processing pipeline for the Pol I-RRN3 and Pol I OC dataset is outlined in Extended Data Fig. 5. First, a small dataset of 967 micrographs was collected on a FEI Talos Arctica. WARP 1.0.7W^73^ was used for the frame alignment, CTF estimation and dose weighting of the micrographs. Using the standard network, 375,673 particles were picked in WARP and extracted into a 220-pixel box. Particles were imported into cryoSPARC and 2D classified. 10 out of 50 classes were selected and 161,124 particles were subjected to *ab initio* reconstruction asking for two classes. One obtained class resembled Pol I only, while the other one, with 45,335 particles, showed additional density next to the stalk where the RRN3 was expected to bind. It was selected and refined with the Homogeneous Refinement tool yielding a map resolving up to 7 Å which was further used as a reference. To obtain a high-resolution data for the Pol I-RRN3-OC sample, we collected a dataset on the Titan Krios TEM. As for Pol I, WARP was used for initial micrograph pre-processing and particle picking. 2,628,144 particles were picked and extracted into a 288-pixel box. The dataset was further processed with RELION 3.1^75^. Micrographs were pre-processed using Relion’s own implementation of MotionCor2 and Gctf, same as for the Pol I EC dataset. Particles were extracted with a 72-pixel box (binned by factor of 4) and split into 5 subsets for faster processing. Each subset was subjected to a global 3D classification with 8 classes using the reference obtained from the FEI Talos Arctica adjusted for the box and pixel size using the EMAN2 package^78^. In each subset, classes containing complete Pol I were selected and those particles were re-extracted into the box of 144 pixels (binned by factor of 2). Using Chimera, the structure of the human Pol I EC was fitted into the map. The structure of the yeast Pol I with RRN3 (PDB: 6RQT)^30^ was matched to it to guide the fitting of the homology model for the human RRN3. A soft mask covering the stalk and the RRN3 was created and it was used for masked 3D classification. In each subset the class that contained a more defined signal in the region of interest was selected. Those particles from all subsets were pooled giving 345,425 particles in total. The resulting intermediate map resolved up to 3.3 Å while data were binned by factor of 2. Another 3D classification with a soft mask on RRN3 only allowed splitting the dataset into two classes: first (referred to as Pol I-RRN3) with 169,513 particles showed signal for the RRN3; and second (referred to as Pol I OC) with 175,912 particles had signal for the nucleic acid, but not for the RRN3. The Pol I-RRN3 class resolved up to 3.4 Å, while the Pol I OC reached 3.6 Å. Both of the classes were unbinned, CTF-refined and polished with the same strategy as for the Pol I EC dataset. The resolution improved up to 3.2 Å for Pol I-RRN3 and to 3.3 Å for Pol I OC, and the obtained maps are denoted as Map D and F respectively. Still, the density corresponding to the upper clamp and the RRN3 in the Map D appeared streaky, possibly due to the clamp movement which distorts the signal. Thus, we applied focused refinement strategy to the intermediate map binned by factor of 2^81^. A soft mask covering the upper clamp and the RRN3 was applied from the 14^th^ iteration of the 3D refinement. The obtained map was used as a reference in 3D classification with a mask covering the stalk and the RRN3. The obtained class with density for RRN3 contained 260,363 particles which were re-extracted into a 288 pixel box. Second round of focused refinement was applied to the unbinned particles in the same manner as previously. The map was then CTF-refined and polished, resulting in a map with the overall resolution of 3.1 Å and improved density for the upper clamp and the RRN3. It was denoted as Map E. Map F corresponding to the Pol I OC was later subjected to the masked classification with a soft mask covering the nucleic acids. A class with improved density corresponding to the DNA scaffold included 164,436 particles and after CTF refinement and polishing, it resolved up to 3.3 Å giving Map G. Maps D, E, F and G were sharpened with LocalDeblur^84^ and LocScale^87^ implemented within CCP-EM^88^.

### Model building and refinement

A homology model based on the yeast Pol I crystal structure (PDB: 4C3I)^19^ was created using: PDB-7AE1^17^ for the subunits shared between Pol I and Pol III; homology models from the SWISS-MODEL Repository^89^ for the subunits RPA1, RPA2, RPA12 and RPA34 and homology models obtained from Phyre2^90^ for RPA43 and RPA49 subunits. Subunits were aligned using UCSF Chimera^80^ and the model was rigid body fitted into Map A. The nucleic acid scaffold from the PDB-6HLR^91^ was fitted into the density within the cleft. The resulting model was further refined using COOT 0.9^92^. First, each chain was fitted as a rigid body into the density and then ProSMART restraints^93^ were generated and used in the refinement to improve the fit. The model was subsequently manually re-built and refined in COOT. To build the downstream DNA, PDB-5M5X^28^ was initially fitted for guidance and then adjusted to fit the cryo-EM density. To build the double-stranded RNA in the RNA exit tunnel, a perfect RNA double-stranded helix (A-form) was fitted and then refined to match the density. The sequence of the nucleic acids was manually mutated to match the used scaffold. Stalk, tWH domain of RPA49 and the RNA in the exit tunnel were built using Map B1. Tracing of the linker of the RPA49 subunit was possible thanks to Map B and was partially aided by the yeast Pol I structure from PDB-6RQT^30^. The structure was iteratively refined using real-space refinement tool^94^ from Phenix 1.13^95^ against Map A with secondary structure restraints turned on. For the two final rounds of the refinement, Map C was used to account for the regions which were more poorly resolved in the Map A.

For building of the Pol I-RRN3 structure, Pol I EC was rigid body fitted into the Map D using Chimera. The nucleic acid scaffold as well as tWH domain of RPA49 were removed. Homology model for the RRN3 was generated using Phyre2 and fitted into the density next to the stalk guided by the structure of yeast Pol I with RRN3 (PDB-6RQT)^30^. Structure for the RPA12 subunit which included the C-terminal domain was taken from the SWISS-MODEL Repository and fitted into the model. In COOT, ProSMART restraints were generated for all subunits and they were refined into the map. For RPA1 and RPA2, subunits were split along the hinges of the clamp to allow more accurate fitting into the map. The upper clamp and active site were extensively re-built manually and refined in COOT. At later stages of the building, Map E was used for the stalk and RRN3 building. Sharpening of the Map D and Map E using LocScale^87^ helped reveal detailed features. Into the extra density inside the DNA binding cleft, a relevant part of the RPA1 subunit homology model from Phyre2 was fitted using Map E and manually adjusted. This section was removed from the final model due to poor quality of the map which did not allow register assignment. The fit of the model into the cryo-EM density was assessed in detail and where necessary it was truncated or extended. The structure was refined against Map E using real-space refinement in Phenix 1.13.

To obtain the model for the Pol I OC, the Pol I-RRN3 structure was fitted into the cryo-EM density. The RRN3 structure was removed. Nucleic acid scaffold from the PDB-5M5W^28^ was rigid body fitted into the visible density inside the cleft using Map G. We truncated the nucleic acids and only left the downstream double-stranded portion of the template. The sequence was manually mutated to match the template used. The structure was manually adjusted and refined in COOT. It was refined against Map F using real-space refinement tool from Phenix 1.13.

Refinement statistics reported within Table 1 were obtained by the MolProbity comprehensive validation tool^96^ implemented within Phenix 1.13. The charge distribution presented in Fig 2. was calculated using the APBS software^97^.

### Data and code availability

Cryo-EM maps obtained within this study have been deposited to the Electron Microscopy Data Bank (EMDB) database under following accession codes: EMD-12795 (Map A, B, B1, B2 and C), EMD-12796 (Map D and E) and EMD-12797 (Map F and G). The atomic models coordinates have been deposited to the Protein Data Bank (PDB) with the following accession codes: 7OB9 (Pol I EC), 7OBA (Pol I-RRN3) and 7OBB (Pol I OC).

Code and further instructions for the phylogenomic analysis are available at: https://github.com/bateman-research/domain-phylo.

## Extended Data Figures and Tables

**Extended Data Fig. 1:**
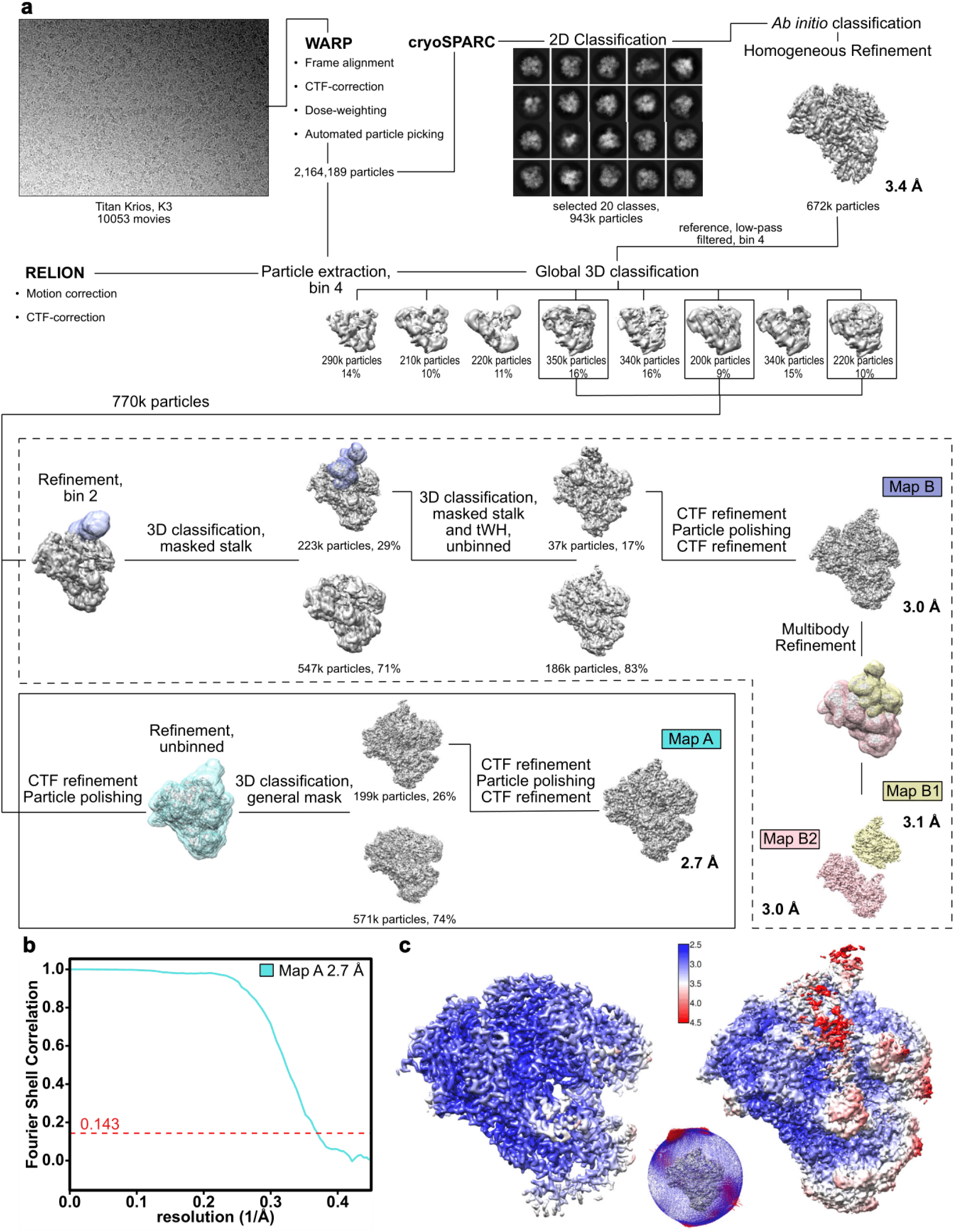
Pol I EC data processing strategy. **a** Processing pipeline for the Pol I EC dataset. Details are described within the STAR Method Details section. Transparent surfaces show masks used for the 3D classification (violet and cyan) or multi-body refinement (yellow and pink)^81^. Shown are unsharpened cryo-EM maps with threshold levels individually adjusted to show relevant features. Reported resolutions were obtained via RELION post-processing. Annotated maps correspond to the maps reported in Table 1 and used for model building. **b** FSC curve for the Pol I EC showing the final average resolution for the map A of 2.7 Å (FSC = 0.143). **c** Local resolution estimation of the Pol I EC obtained with the script implemented in RELION. The map at a higher threshold (left) shows the high resolution core, while at lower threshold values (right), lower resolution peripheral subunits become visible. Angular distribution plot of all particles contributing to the Pol I EC structure is shown next to the local resolution maps.

**Extended Data Fig. 2:**
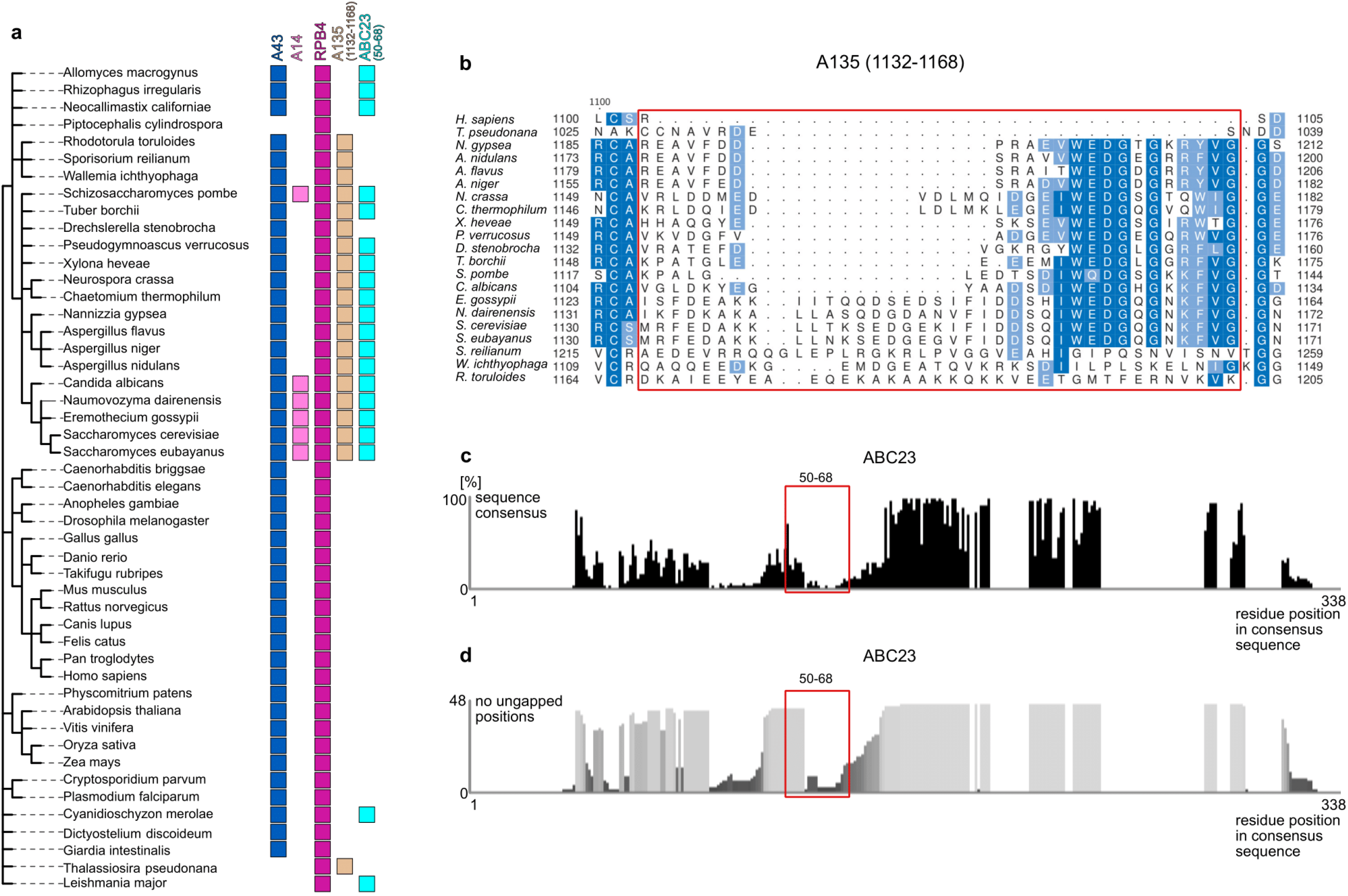
Phylogenomic analysis of the stalk subunits and extensions from A135 and ABC23 contacting the stalk. **a** Analysis of A14 and A43 Pol I stalk subunits, smaller stalk subunit RPB4 from Pol II and extensions from subunits A135 (1132-1168) and ABC23 (50-68) as indicated in Fig. 2. Filled square indicates presence of a homologous protein in the corresponding species. Color code as in Fig. 3. Genes are annotated per *S. cerevisiae* nomenclature. For UniProt identifiers of the identified homologues see Supplementary Table 2. **b** Sequence alignment of subunit A135 across species where the *S. cerevisiae* 1132-1168 insert was identified. Human sequence of RPA2 subunit is included in top row for comparison. Sequences are sorted by alignment score. **c** Sequence consensus score plotted across the length of the consensus sequence for the ABC23 subunit. Percentage of sequences with the consensus residue is shown. **d** Occupancy plot for the ABC23 subunit showing the number of ungapped residues in each position of the consensus alignment. Total of 48 sequences were aligned. **b-d** The red box marks the location of the inserts of interest. Numbering in reference to *S. cerevisiae* sequence.

**Extended Data Fig. 3:**
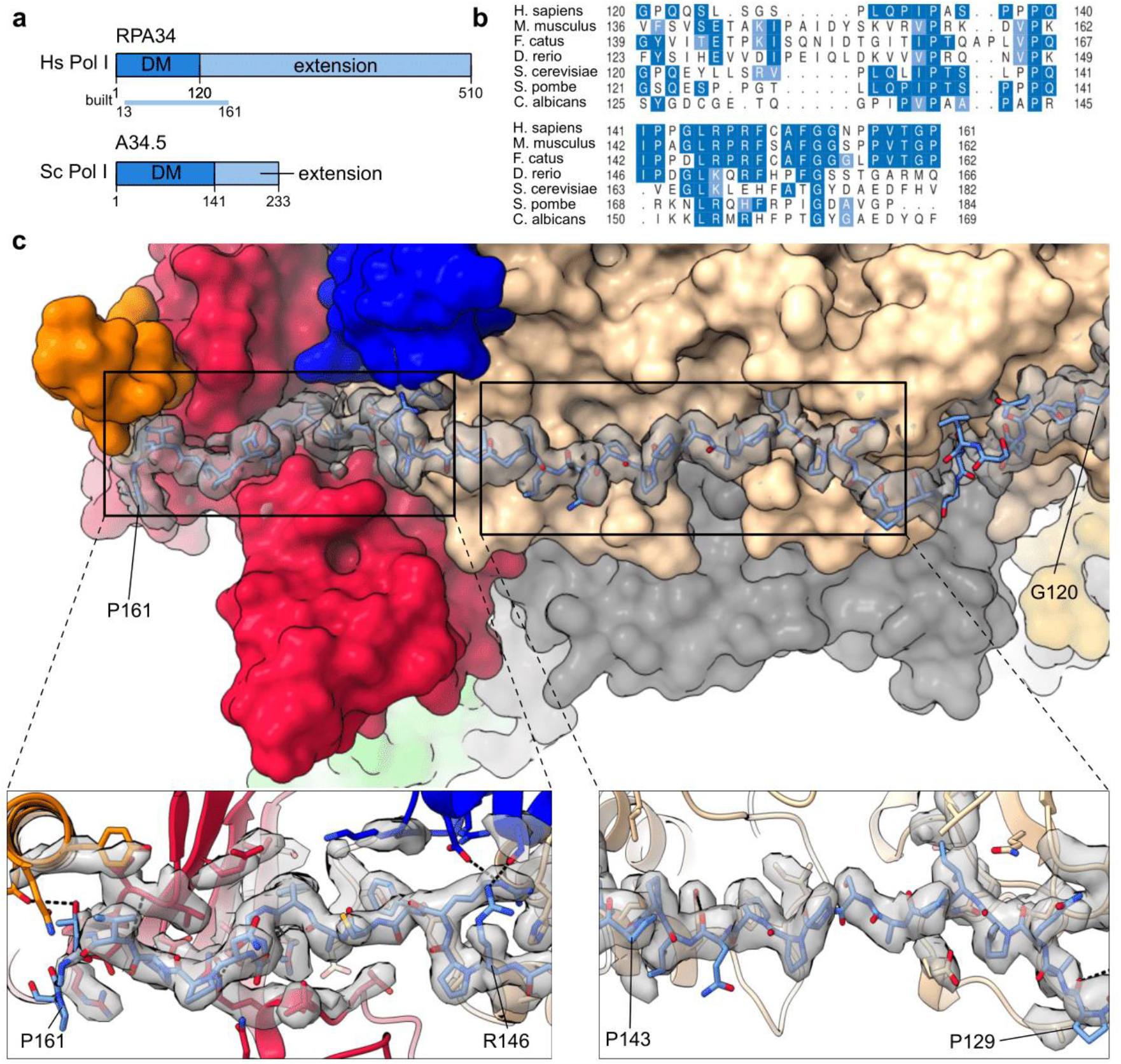
The RPA34 extension tethers the heterodimer to the Pol I core. **a** Cartoon domain representation of the domain composition of the human RPA34 (top) and its yeast homologue A34.5 (bottom). The colored bar indicates the region modelled in the structure denoted as ‘built’. DM – dimerization domain. **b** Multiple sequence alignment of the modelled portion of the RPA34 extension between different species listed on the left. Dark blue marks identical residues and light blue similar residues. Dots mark gaps in the sequence. **c** RPA34 extension path (stick representation) on the Pol I surface (surface representation). Cryo-EM density, corresponding to the extension is shown in grey, transparent representation. Lower panels show the close up view of the extension (stick representation) surrounded by the Pol I core (cartoon representation). Residues from the Pol I core within the 5 Å radius from the linker are shown as sticks. Putative hydrogen bonds are shown as black dotted lines.

**Extended Data Fig. 4:**
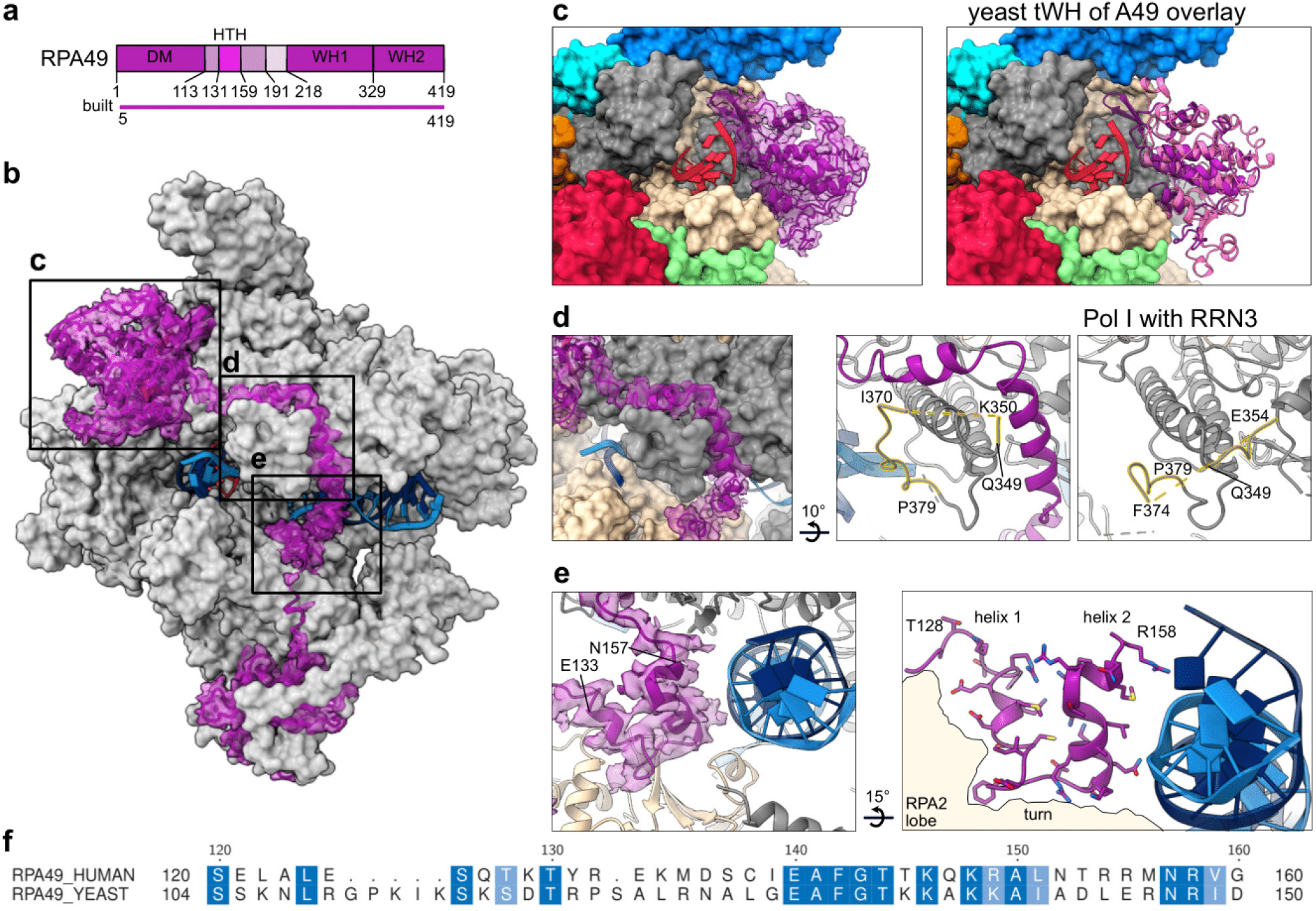
RPA49 linker runs along the Pol I clamp to position the tWH domain. **a** Cartoon domain representation of the RPA49 subunit. The colored bar indicates the region modelled in the structure denoted as ‘built’. DM – dimerization domain; HTH – helix-turn-helix motif; WH – winged helix domain. **b** Overall arrangement of the RPA49 subunit (purple, cartoon representation fitted into the cryo-EM density extracted from Map C) on the surface of Pol I (grey, surface representation). The DNA scaffold is shown as a cartoon. **c** Positioning of the tWH domain (purple, cartoon representation fitted into the corresponding cryo-EM density) near the RNA exit tunnel. The Pol I core is shown in a surface representation colored by subunit. Right panel: yeast tWH domain (pink, cartoon representation) from PDB: 5M64^28^ overlaid onto the human Pol I structure. **d** Left panel: the linker (purple) wraps around a knob formed by clamp coiled-coils (grey, surface representation). Middle panel: partially disordered loop (gold highlight) connecting the coiled-coils (grey cartoon representation) allows binding of the RPA49 linker (purple cartoon). Right panel: in the structure of human Pol I bound to RRN3, where the RPA49 linker is disordered, the same loop (golden highlight) adopts a conformation that would clash with the RPA49 linker. **e** HTH motif within the linker (purple with the corresponding cryo-EM density shown in transparent representation) is positioned close to the downstream DNA helix (blue, cartoon representation). Right panel: side chains of the HTH are shown in stick representation (purple). Helix 1 and turn contact the RPA2 lobe (wheat outline), while helix 2 can contact the downstream DNA. **f** Sequence alignment of the part of RPA49 linker containing the HTH motif between human and yeast. Dark blue marks identical residues and light blue similar residues. Dots mark gaps in the sequence.

**Extended Data Fig. 5:**
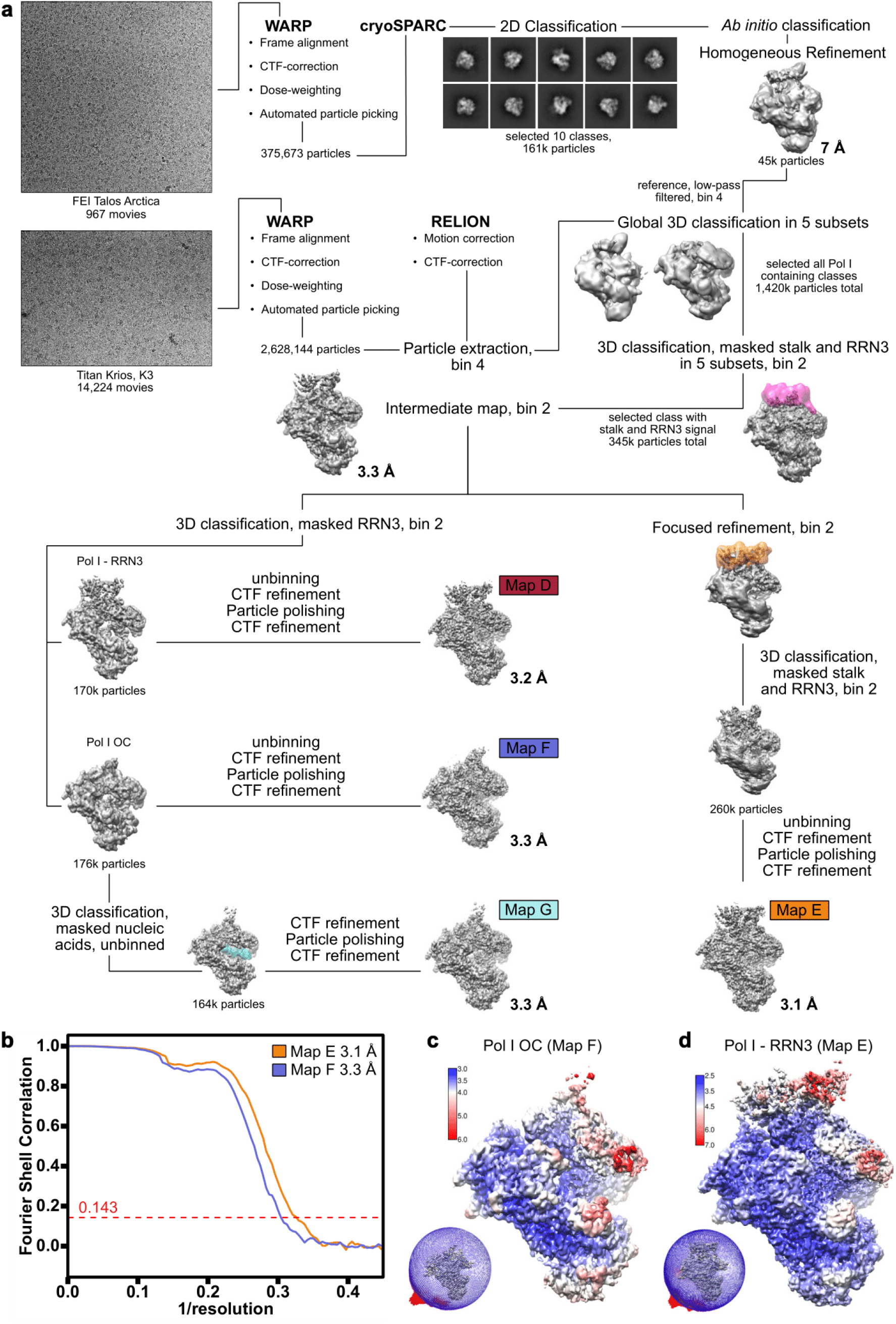
Pol I-RRN3 and Pol I OC dataset processing strategy. **a** Details of the pipeline are described within the STAR Method Details section. A small initial dataset was collected using FEI Talos Arctica and processed in cryoSPARC to obtain a reference that was used for global 3D classification for the high-resolution dataset collected on Titan Krios. First global 3D classification was performed with the dataset split into 5 subsets and representative selected classes are shown. Transparent surfaces show masks used for the 3D classification (pink and cyan) or for focused refinement (orange). Shown are unsharpened cryo-EM maps with threshold level individually adjusted to show relevant features. Reported resolutions were obtained via RELION post-processing. Annotated maps correspond to the maps reported in Table 1 and used for model building. **b** FSC curves for the Pol I-RRN3 and Pol I OC showing the final average resolution for the maps E and F of 3.1 and 3.3 Å respectively (FSC = 0.143). **c, d** Local resolution estimation of Pol I OC (**c**) and Pol I-RRN3 (**d**) obtained with the RELION implementation. Maps are shown at lower threshold values, so that the lower resolution peripheral subunits become visible. Angular distribution plot of all particles contributing to the structures is shown to the left of each local resolution map.

**Extended Data Fig. 6.**
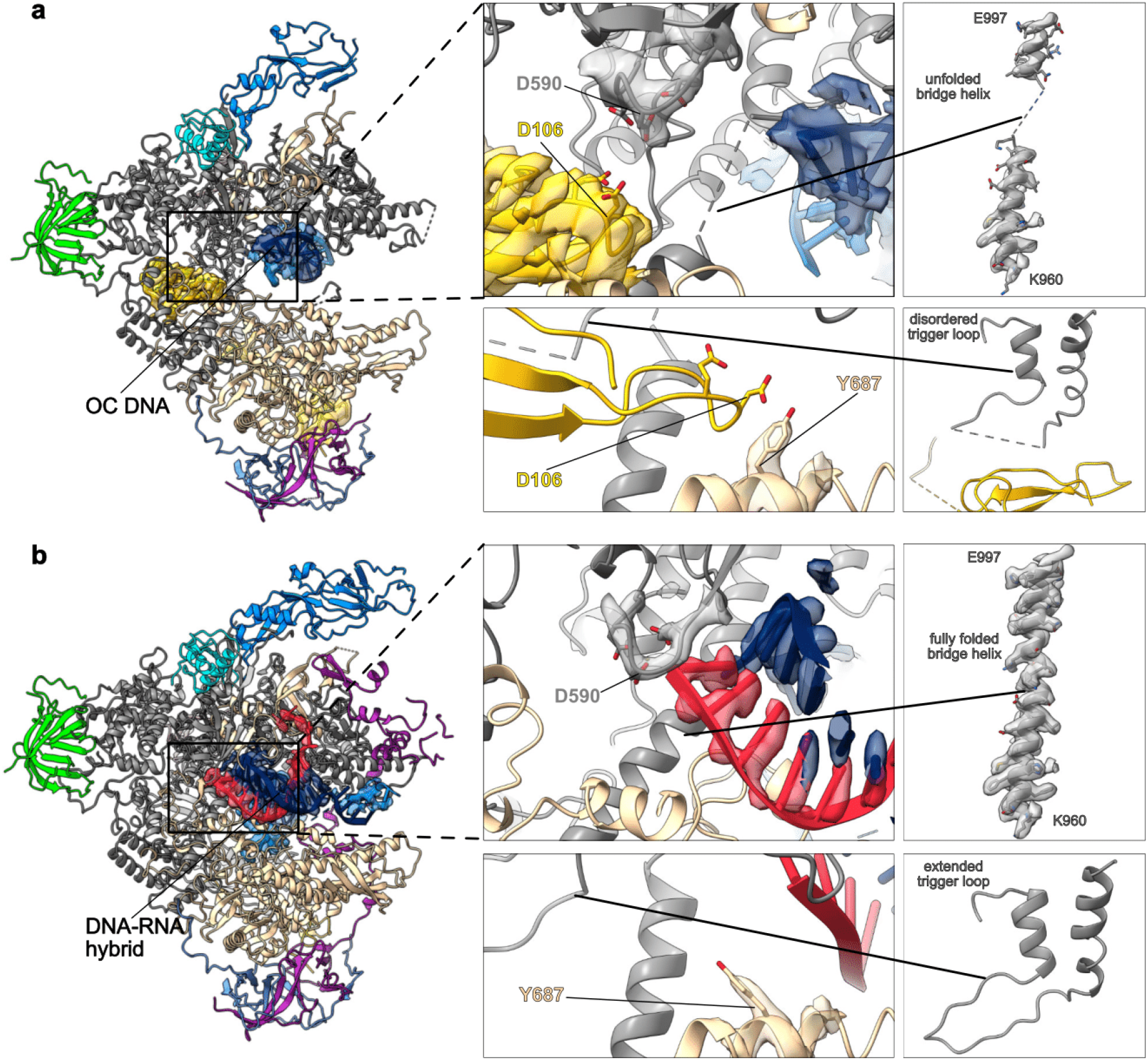
Pol I OC represents an inactive state of Pol I. **a, b** Pol I OC (**a**) and Pol I EC (**b**) in cartoon representation with subunits colored as in Fig. 1. The extracted cryo-EM density for the relevant parts of the structures is shown in transparent representation colored according to the respective subunit. Subunits directly occluding the view of the active site are hidden. **a** Insertion of the RPA12 C-terminal domain (yellow) into the active site. Aspartates in the RPA12 catalytic loop are shown in stick representation. **a, b left panels** Structures overview. **a, b top middle panels** Catalytic triad of the aspartates (D588, D590 and D592) is shown in stick representation. **a, b bottom middle panels** Conformation of the gating tyrosine (Y687) changes between the Pol I OC and Pol I EC. **a, b right panels** Functional elements of the active site such as bridge helix and trigger loop are disordered in Pol I OC (**a right panels**), while in the Pol I EC (**b right panels**) they are fully folded.

**Extended Data Fig. 7:**
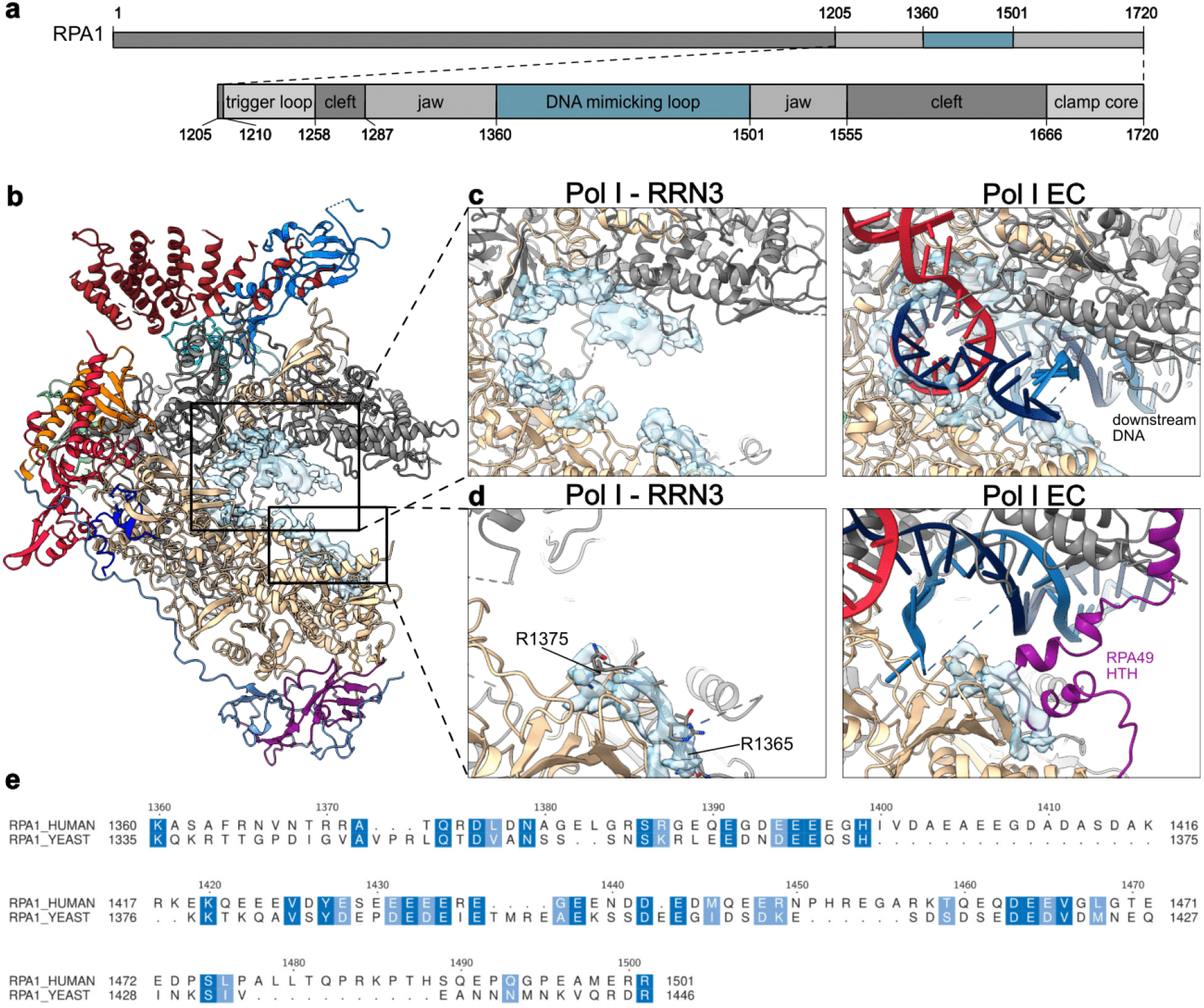
Density corresponding to the DNA mimicking loop in the Pol I-RRN3 map. **a** Cartoon domain representation of the RPA1 subunit. The C-terminus harboring the jaw extension termed ‘DNA-mimicking loop’ is shown on the close-up view. **b-d** Structures are shown in cartoon representation. Additional density lining the DNA binding cleft in the Pol I-RRN3 structure which likely corresponds to the DNA-mimicking loop is shown as transparent, blue surface. **b** Overview representation of the Pol I-RRN3 structure. **c** Close-up view of the additional density in the DNA binding cleft (left panel), which is then overlaid on the Pol I EC structure (right panel), where it would overlap with the DNA backbone. **d** Close-up view of the first 10-residues of the DNA mimicking loop (stick representation) which bind to the RPA2 core (left panel). When the cryo-EM density corresponding to those residues is superimposed on the Pol I EC (right panel) it overlaps with the turn from the HTH motif of subunit RPA49 (purple).

**Supplementary Table 1:** Mass spectrometry assessment of the purified Pol I complex. Statistics for the highest scoring hit used for the identification of the subunit present in the protein band as visualized by coomassie stain in Fig. 1b.

| Pol I subunit | Gene name | Number of unique peptides | Sequence coverage [%] | Max score |
| --- | --- | --- | --- | --- |
| RPA1 | *POLR1A* | 246 | 89.9 | 222 |
| RPA2 | *POLR1B* | 93 | 51.8 | 121 |
| RPAC1 | *POLR1C* | 38 | 79.9 | 177 |
| RPA49 | *POLR1E* | 55 | 67.4 | 137 |
| RPA43 | *TWISTNB* | 4 | 18.0 | 95 |
| RPA34 | *CD3EAP* | 8 | 10.8 | 70 |
| RPABC1 | *POLR2E* | 40 | 79.6 | 144 |
| RPABC3 | *POLR2H* | 20 | 78.8 | 183 |
| RPABC2 | *POLR2F* | 4 | 14.8 | 71 |
| RPA12 | *ZNRD1* | 14 | 63.0 | 82 |
| RPAC2 | *POLR1D* | 16 | 63.4 | 102 |
| RPABC4 | *POLR2K* | 5 | 45.8 | 82 |
| RPABC5 | *POLR2L* | 2 | 29.4 | 64 |

**Supplementary Table 2:**
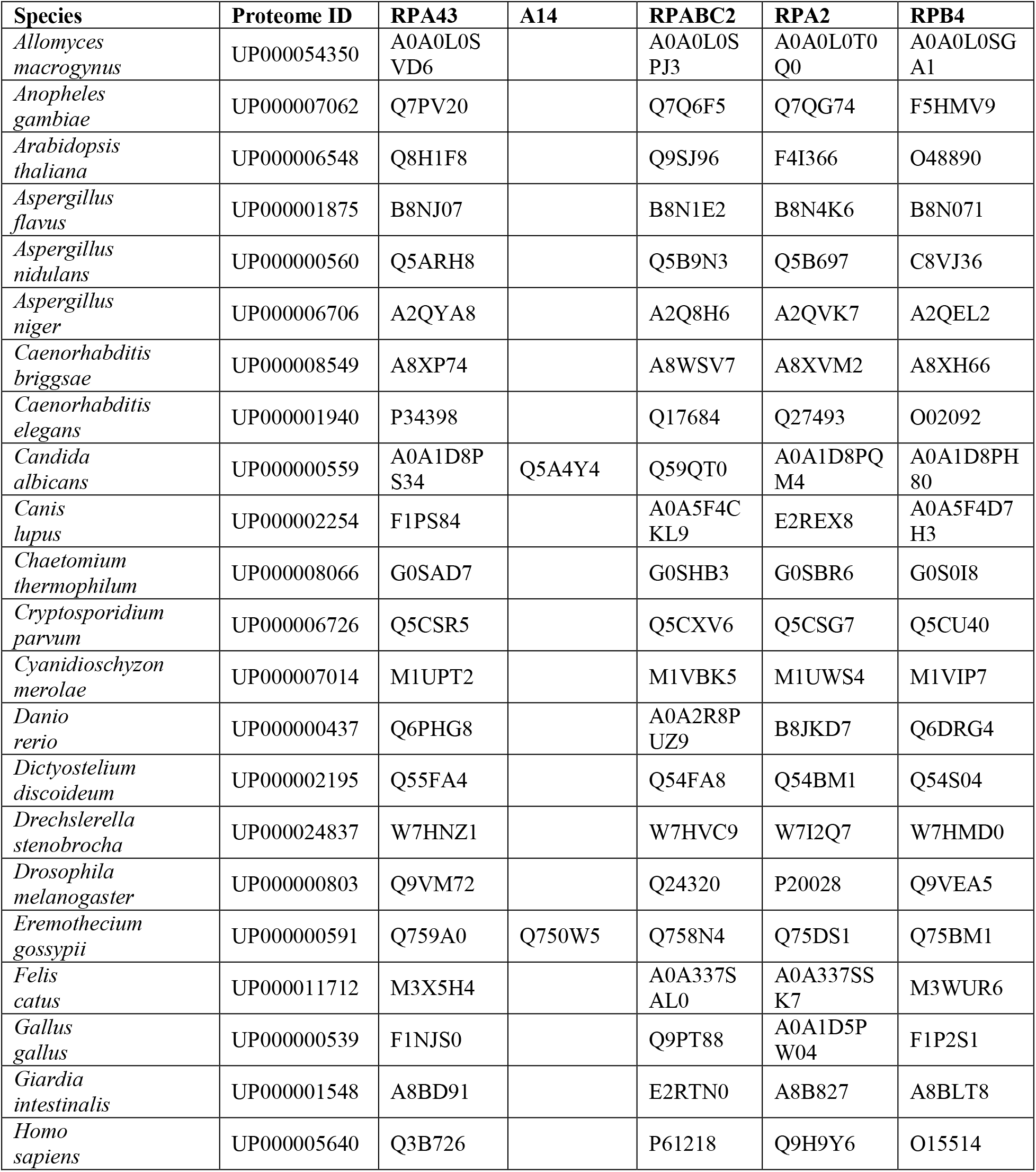
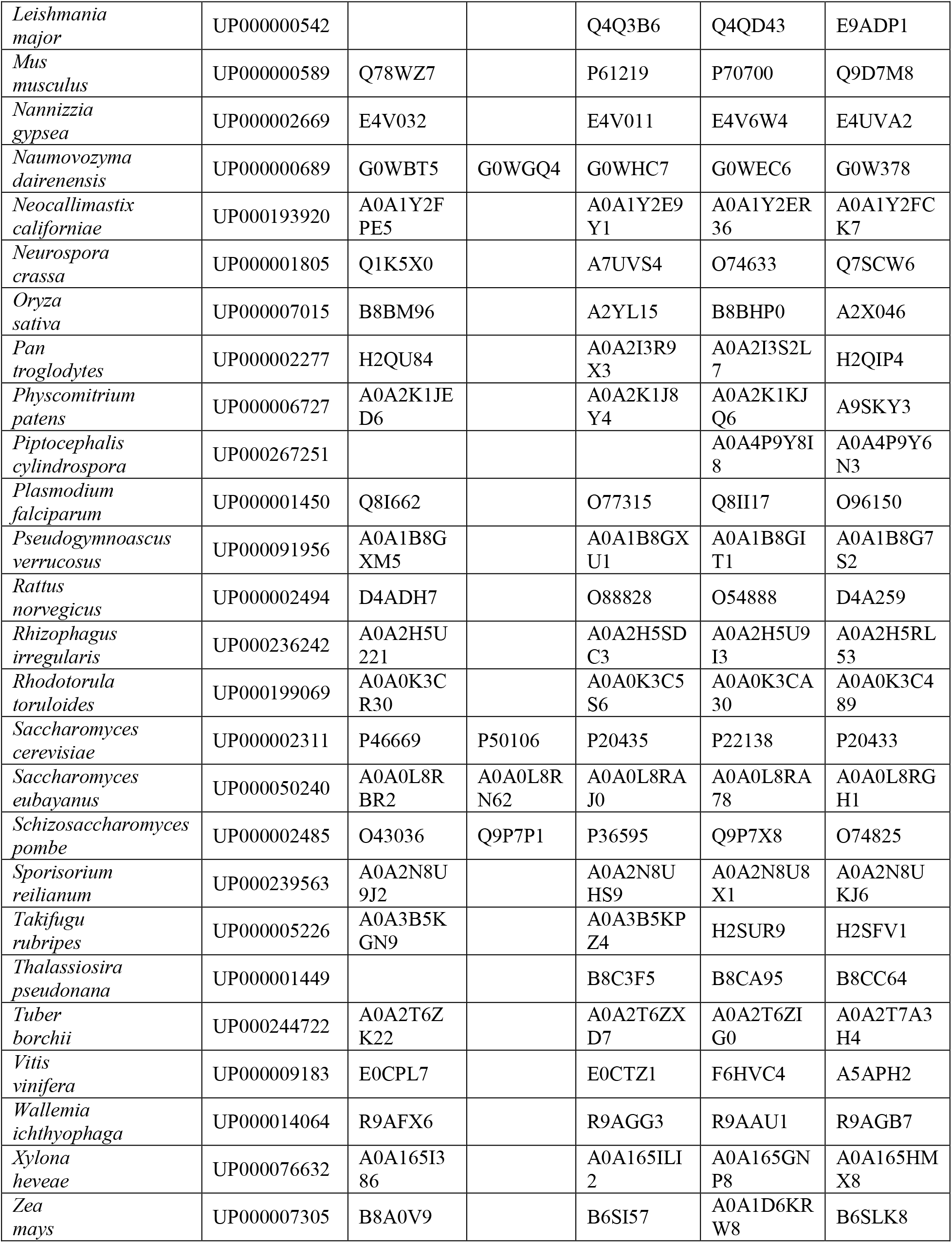
Phylogenomic analysis of the stalk subunits and core contacting the stalk. Listed are analyzed species, their proteome identifiers and UniProt identifiers for homologs of RPA43, A14 (*S. cerevisiae* smaller stalk subunit), RPABC2, RPA2 and RPB4 (Pol II smaller stalk subunit). Homologs were retrieved with the HMMER tool ^66^ as described in the Methods details. See Extended Data Fig. 2A for the corresponding phylogenomic tree.

## References

1. Goodfellow, S. J. & Zomerdijk, J. C. B. M. Basic mechanisms in RNA polymerase I transcription of the ribosomal RNA genes. Subcell. Biochem. 61, 211–236 (2013).

2. Palazzo, A. F. & Lee, E. S. Non-coding RNA: What is functional and what is junk? Front. Genet. 5, 2 (2015).

3. Ferreira, R., Schneekloth, J. S., Panov, K. I., Hannan, K. M. & Hannan, R. D. Targeting the RNA polymerase I transcription for cancer therapy comes of age. Cells 9, 266 (2020).

4. Bywater, M. J. et al. Inhibition of RNA polymerase I as a therapeutic strategy to promote cancer-specific activation of p53. Cancer Cell 22, 51–65 (2012).

5. Sharifi, S. & Bierhoff, H. Regulation of RNA polymerase I transcription in development, disease, and aging. Annu. Rev. Biochem. 87, 51–73 (2018).

6. Martínez Corrales, G. et al. Partial Inhibition of RNA Polymerase I Promotes Animal Health and Longevity. Cell Rep. 30, 1661–1669.e4 (2020).

7. Hannan, K. M., Sanij, E., Rothblum, L. I., Hannan, R. D. & Pearson, R. B. Dysregulation of RNA polymerase I transcription during disease. Biochim. Biophys. Acta 1829, 342–360 (2013).

8. Weaver, K. N. et al. Acrofacial Dysostosis, Cincinnati type, a Mandibulofacial Dysostosis syndrome with limb anomalies, is caused by POLR1A dysfunction. Am. J. Hum. Genet. 96, 765–774 (2015).

9. Sanchez, E. et al. POLR1B and neural crest cell anomalies in Treacher Collins syndrome type 4. Genet. Med. 22, 547–556 (2020).

10. Matsumoto, N. et al. Treacher Collins syndrome 3 (TCS3)-associated POLR1C mutants are localized in the lysosome and inhibits chondrogenic differentiation. Biochem. Biophys. Res. Commun. 499, 78–85 (2018).

11. Vincent, M. et al. Treacher Collins syndrome: A clinical and molecular study based on a large series of patients. Genet. Med. 18, 49–56 (2016).

12. Gauquelin, L. et al. Clinical spectrum of POLR3-related leukodystrophy caused by biallelic POLR1C pathogenic variants. Neurol. Genet. 5, e369 (2019).

13. Vannini, A. & Cramer, P. Conservation between the RNA polymerase I, II, and III transcription initiation machineries. Mol. Cell 45, 439–446 (2012).

14. Khatter, H., Vorländer, M. K. & Müller, C. W. RNA polymerase I and III: similar yet unique. Curr. Opin. Struct. Biol. 47, 88–94 (2017).

15. Cramer, P., Bushnell, D. A. & Kornberg, R. D. Structural basis of transcription: RNA polymerase II at 2.8 ångstrom resolution. Science 292, 1863–1876 (2001).

16. Bernecky, C., Herzog, F., Baumeister, W., Plitzko, J. M. & Cramer, P. Structure of transcribing mammalian RNA polymerase II. Nature 529, 551–554 (2016).

17. Girbig, M. et al. Cryo-EM structures of human RNA polymerase III in its unbound and transcribing states. Nat. Struct. Mol. Biol. 28, 210–219 (2021).

18. Hoffmann, N. A. et al. Molecular structures of unbound and transcribing RNA polymerase III. Nature 528, 231–236 (2015).

19. Fernández-Tornero, C. et al. Crystal structure of the 14-subunit RNA polymerase I. Nature 502, 644–649 (2013).

20. Engel, C., Sainsbury, S., Cheung, A. C., Kostrewa, D. & Cramer, P. RNA polymerase I structure and transcription regulation. Nature 502, 650–655 (2013).

21. Russell, J. & Zomerdijk, J. C. B. M. The RNA polymerase I transcription machinery. Biochem. Soc. Symp. 73, 203–216 (2006).

22. Grummt, I. Life on a planet of its own: regulation of RNA polymerase I transcription in the nucleolus. Genes Dev. 17, 1691–1702 (2003).

23. Heix, J. et al. Mitotic silencing of human rRNA synthesis: Inactivation of the promoter selectivity factor SL1 by cdc2/cyclin B-mediated phosphorylation. EMBO J. 17, 7373–7381 (1998).

24. Chen, S. et al. Repression of RNA polymerase I upon stress is caused by inhibition of RNA-dependent deacetylation of PAF53 by SIRT7. Mol. Cell 52, 303–313 (2013).

25. Mayer, C., Bierhoff, H. & Grummt, I. The nucleolus as a stress sensor: JNK2 inactivates the transcription factor TIF-IA and down-regulates rRNA synthesis. Genes Dev. 19, 933– 941 (2005).

26. Mayer, C., Zhao, J., Yuan, X. & Grummt, I. mTOR-dependent activation of the transcription factor TIF-IA links rRNA synthesis to nutrient availability. Genes Dev. 18, 423–434 (2004).

27. Zhao, J., Yuan, X., Frödin, M. & Grummt, I. ERK-dependent phosphorylation of the transcription initiation factor TIF-IA is required for RNA polymerase I transcription and cell growth. Mol. Cell 11, 405–413 (2003).

28. Tafur, L. et al. Molecular Structures of Transcribing RNA Polymerase I. Mol. Cell 64, 1135–1143 (2016).

29. Sadian, Y. et al. Structural insights into transcription initiation by yeast RNA polymerase I. EMBO J. 36, 2698–2709 (2017).

30. Sadian, Y. et al. Molecular insight into RNA polymerase I promoter recognition and promoter melting. Nat. Commun. 10, 1–13 (2019).

31. Han, Y. et al. Structural mechanism of ATP-independent transcription initiation by RNA polymerase I. Elife 6, e27414 (2017).

32. Engel, C., Plitzko, J. & Cramer, P. RNA polymerase I-Rrn3 complex at 4.8 Å resolution. Nat. Commun. 7, 1–5 (2016).

33. Neyer, S. et al. Structure of RNA polymerase I transcribing ribosomal DNA genes. Nature 540, 607–610 (2016).

34. Pilsl, M. & Engel, C. Structural basis of RNA polymerase I pre-initiation complex formation and promoter melting. Nat. Commun. 11, 1–10 (2020).

35. Engel, C. et al. Structural Basis of RNA Polymerase I Transcription Initiation. Cell 169, 120–131.e22 (2017).

36. Geiger, S. R. et al. RNA polymerase I contains a TFIIF-related DNA-binding subcomplex. Mol. Cell 39, 583–594 (2010).

37. Knutson, B. A., McNamar, R. & Rothblum, L. I. Dynamics of the RNA polymerase I TFIIF/TFIIE-like subcomplex: A mini-review. Biochem. Soc. Trans. 48, 1917–1927 (2020).

38. Wang, Q. et al. Structural insights into transcriptional regulation of human RNA polymerase III. Nat. Struct. Mol. Biol. 28, 220–227 (2021).

39. Kang, J. Y. et al. RNA polymerase accommodates a pause RNA hairpin by global conformational rearrangements that prolong pausing. Mol. Cell 69, 802–815.e1 (2018).

40. Turowski, T. W. et al. Nascent transcript folding plays a major role in determining RNA polymerase elongation rates. Mol. Cell 79, 488–503.e11 (2020).

41. Fromont-Racine, M., Senger, B., Saveanu, C. & Fasiolo, F. Ribosome assembly in eukaryotes. Gene 313, 17–42 (2003).

42. Heiss, F. B., Daiß, J. L., Becker, P. & Engel, C. Conserved strategies of RNA polymerase I hibernation and activation. Nat. Commun. 12, 1–9 (2021).

43. He, Y. et al. Near-atomic resolution visualization of human transcription promoter opening. Nature 533, 359–365 (2016).

44. Kay, B. K., Williamson, M. P. & Sudol, M. The importance of being proline: the interaction of proline-rich motifs in signaling proteins with their cognate domains. FASEB J. 14, 231– 241 (2000).

45. Albert, B. et al. RNA polymerase I-specific subunits promote polymerase clustering to enhance the rRNA gene transcription cycle. J. Cell Biol. 192, 277–293 (2011).

46. Yamamoto, K. et al. Multiple protein-protein interactions by RNA polymerase I-associated factor PAF49 and role of PAF49 in rRNA transcription. Mol. Cell. Biol. 24, 6338–6349 (2004).

47. Penrod, Y., Rothblum, K., Cavanaugh, A. & Rothblum, L. I. Regulation of the association of the PAF53/PAF49 heterodimer with RNA polymerase I. Gene 556, 61–67 (2015).

48. Panov, K. I. et al. RNA polymerase I-specific subunit CAST/hPAF49 has a role in the activation of transcription by UpstreamBinding Factor. Mol. Cell. Biol. 26, 5436–5448 (2006).

49. McNamar, R., Abu-Adas, Z., Rothblum, K., Knutson, B. A. & Rothblum, L. I. Conditional depletion of the RNA polymerase I subunit PAF53 reveals that it is essential for mitosis and enables identification of functional domains. J. Biol. Chem. 294, 19907–19922 (2019).

50. Moorefield, B., Greene, E. A. & Reeder, R. H. RNA polymerase I transcription factor Rrn3 is functionally conserved between yeast and human. Proc. Natl. Acad. Sci. U. S. A. 97, 4724–4729 (2000).

51. Miller, G. et al. hRRN3 is essential in the SL1-mediated recruitment of RNA polymerase I to rRNA gene promoters. EMBO J. 20, 1373–1382 (2001).

52. Darrière, T. et al. Genetic analyses led to the discovery of a super-active mutant of the RNA polymerase I. PLOS Genet. 15, e1008157 (2019).

53. Ruan, W., Lehmann, E., Thomm, M., Kostrewa, D. & Cramer, P. Evolution of two modes of intrinsic RNA polymerase transcript cleavage. J. Biol. Chem. 286, 18701–18707 (2011).

54. Cheung, A. C. M. & Cramer, P. Structural basis of RNA polymerase II backtracking, arrest and reactivation. Nature 471, 249–253 (2011).

55. Yuan, X., Zhao, J., Zentgraf, H., Hoffmann-Rohrer, U. & Grummt, I. Multiple interactions between RNA polymerase I, TIF-IA and TAF _I_ subunits regulate preinitiation complex assembly at the ribosomal gene promoter. EMBO Rep. 3, 1082–1087 (2002).

56. Bierhoff, H., Dundr, M., Michels, A. A. & Grummt, I. Phosphorylation by Casein Kinase 2 facilitates rRNA gene transcription by promoting dissociation of TIF-IA from elongating RNA polymerase I. Mol. Cell. Biol. 28, 4988–4998 (2008).

57. Dephoure, N. et al. A quantitative atlas of mitotic phosphorylation. Proc. Natl. Acad. Sci. 105, 10762–10767 (2008).

58. Cavanaugh, A. H. et al. Rrn3 phosphorylation is a regulatory checkpoint for ribosome biogenesis. J. Biol. Chem. 277, 27423–27432 (2002).

59. Schaefer, E. et al. Autosomal recessive POLR1D mutation with decrease of TCOF1 mRNA is responsible for Treacher Collins syndrome. Genet. Med. 16, 720–724 (2014).

60. Dauwerse, J. G. et al. Mutations in genes encoding subunits of RNA polymerases I and III cause Treacher Collins syndrome. Nat. Genet. 43, 20–22 (2011).

61. Thiffault, I. et al. Recessive mutations in POLR1C cause a leukodystrophy by impairing biogenesis of RNA polymerase III. Nat. Commun. 6, 25 (2015).

## References (Methods)

62. Ran, F. A. et al. Genome engineering using the CRISPR-Cas9 system. Nat. Protoc. 8, 2281– 2308 (2013).

63. Schindelin, J. et al. Fiji: An open-source platform for biological-image analysis. Nat. Methods 9, 676–682 (2012).

64. Bateman, A. et al. UniProt: The universal protein knowledgebase in 2021. Nucleic Acids Res. 49, D480–D489 (2021).

65. Federhen, S. The NCBI Taxonomy database. Nucleic Acids Res. 40, D136–D143 (2012).

66. Eddy, S. R. Accelerated profile HMM searches. PLoS Comput. Biol. 7, 1002195 (2011).

67. Mistry, J. et al. Pfam: The protein families database in 2021. Nucleic Acids Res. 49, D412– D419 (2021).

68. Edgar, R. C. MUSCLE: Multiple sequence alignment with high accuracy and high throughput. Nucleic Acids Res. 32, 1792–1797 (2004).

69. Letunic, I. & Bork, P. Interactive Tree of Life (iTOL) v4: Recent updates and new developments. Nucleic Acids Res. 47, W256–W259 (2019).

70. Waterhouse, A. M., Procter, J. B., Martin, D. M. A., Clamp, M. & Barton, G. J. Jalview Version 2 - a multiple sequence alignment editor and analysis workbench. Bioinformatics 25, 1189–1191 (2009).

71. Madeira, F. et al. The EMBL-EBI search and sequence analysis tools APIs in 2019. Nucleic Acids Res. 47, W636–W641 (2019).

72. Mastronarde, D. N. Automated electron microscope tomography using robust prediction of specimen movements. J. Struct. Biol. 152, 36–51 (2005).

73. Tegunov, D. & Cramer, P. Real-time cryo-electron microscopy data preprocessing with Warp. Nat. Methods 16, 1146–1152 (2019).

74. Punjani, A., Rubinstein, J. L., Fleet, D. J. & Brubaker, M. A. CryoSPARC: Algorithms for rapid unsupervised cryo-EM structure determination. Nat. Methods 14, 290–296 (2017).

75. Zivanov, J. et al. New tools for automated high-resolution cryo-EM structure determination in RELION-3. Elife 7, e42166 (2018).

76. Zheng, S. Q. et al. MotionCor2: Anisotropic correction of beam-induced motion for improved cryo-electron microscopy. Nat. Methods 14, 331–332 (2017).

77. Zhang, K. Gctf: Real-time CTF determination and correction. J. Struct. Biol. 193, 1–12 (2016).

78. Tang, G. et al. EMAN2: An extensible image processing suite for electron microscopy. J. Struct. Biol. 157, 38–46 (2007).

79. Zivanov, J., Nakane, T. & Scheres, S. H. W. Estimation of high-order aberrations and anisotropic magnification from cryo-EM data sets in RELION-3.1. IUCrJ 7, 253–267 (2020).

80. Pettersen, E. F. et al. UCSF Chimera - A visualization system for exploratory research and analysis. J. Comput. Chem. 25, 1605–1612 (2004).

81. Nakane, T., Kimanius, D., Lindahl, E. & Scheres, S. H. W. Characterisation of molecular motions in cryo-EM single-particle data by multi-body refinement in RELION. Elife 7, e36861 (2018).

82. Rosenthal, P. B. & Henderson, R. Optimal determination of particle orientation, absolute hand, and contrast loss in single-particle electron cryomicroscopy. J. Mol. Biol. 333, 721– 745 (2003).

83. Liebschner, D. et al. Macromolecular structure determination using X-rays, neutrons and electrons: Recent developments in Phenix. Acta Crystallogr. Sect. D 75, 861–877 (2019).

84. Ramírez-Aportela, E. et al. Automatic local resolution-based sharpening of cryo-EM maps. Bioinformatics 36, 765–772 (2019).

85. de la Rosa-Trevín, J. M. et al. Scipion: A software framework toward integration, reproducibility and validation in 3D electron microscopy. J. Struct. Biol. 195, 93–99 (2016).

86. Terwilliger, T. C., Sobolev, O. V., Afonine, P. V. & Adams, P. D. Automated map sharpening by maximization of detail and connectivity. Acta Crystallogr. Sect. D 74, 545– 559 (2018).

87. Jakobi, A. J., Wilmanns, M. & Sachse, C. Model-based local density sharpening of cryo-EM maps. Elife 6, e27131 (2017).

88. Burnley, T., Palmer, C. M. & Winn, M. Recent developments in the CCP-EM software suite. in *Acta Crystallogr*. Sect. D 73, 469–477 (2017).

89. Bienert, S. et al. The SWISS-MODEL Repository - New features and functionality. Nucleic Acids Res. 45, D313–D319 (2017).

90. Kelley, L. A., Mezulis, S., Yates, C. M., Wass, M. N. & Sternberg, M. J. E. The Phyre2 web portal for protein modeling, prediction and analysis. Nat. Protoc. 10, 845–858 (2015).

91. Tafur, L. et al. The cryo-EM structure of a 12-subunit variant of RNA polymerase I reveals dissociation of the A49-A34.5 heterodimer and rearrangement of subunit A12.2. Elife 8, (2019).

92. Casañal, A., Lohkamp, B. & Emsley, P. Current developments in *Coot* for macromolecular model building of Electron Cryo-microscopy and Crystallographic Data. Protein Sci. 29, 1055–1064 (2020).

93. Nicholls, R. A., Fischer, M., Mcnicholas, S. & Murshudov, G. N. Conformation-independent structural comparison of macromolecules with ProSMART. Acta Crystallogr. Sect. D Biol. Crystallogr. 70, 2487–2499 (2014).

94. Afonine, P. V. et al. Real-space refinement in PHENIX for cryo-EM and crystallography. Acta Crystallogr. Sect. D Struct. Biol. 74, 531–544 (2018).

95. Adams, P. D. et al. PHENIX: A comprehensive Python-based system for macromolecular structure solution. Acta Crystallogr. Sect. D Biol. Crystallogr. 66, 213–221 (2010).

96. Davis, I. W. et al. MolProbity: All-atom contacts and structure validation for proteins and nucleic acids. Nucleic Acids Res. 35, (2007).

97. Jurrus, E. et al. Improvements to the APBS biomolecular solvation software suite. Protein Sci. 27, 112–128 (2018).

